# Improving eDNA filtration and purification for qPCR detection of *Schistosoma mansoni* in lakes Albert and Victoria

**DOI:** 10.1101/2025.03.30.646195

**Authors:** Cecilia Wangari Wambui, Mila Viaene, Hannah Njiriku Mwangi, Benjamin André, David Were Oguttu, Casim Umba Tolo, Bart Hellemans, Tine Huyse, Hugo F. Gante

**Author notes:** Contributed equally to this work: Cecilia Wangari Wambui and Mila Viaene.

## Abstract

**Background:** Schistosomiasis, a parasitic disease caused by *Schistosoma* trematodes, remains a significant public health burden in sub-Saharan Africa, particularly in regions with limited access to clean water, sanitation and hygiene. Effective disease control strategies rely on large-scale surveillance to accurately identify and target high-risk populations. However, traditional methods such as malacological surveys and stool/urine microscopy in humans often lack sensitivity and scalability. Environmental DNA (eDNA) is emerging as a promising tool for non-invasive surveillance of aquatic pathogens, offering enhanced sensitivity and feasibility for large-scale monitoring.

**Methods:** This study evaluated the efficacy of eDNA-based detection of *Schistosoma mansoni* in water samples from Lake Albert and Lake Victoria, Uganda. Three eDNA filtration techniques: open membrane, Waterra capsule and Sylphium capsule filters, were compared for DNA yield and detection efficiency. SYBR Green quantitative polymerase chain reaction (qPCR) targeting the mitochondrial cytochrome oxidase subunit I (COI) gene was performed to quantify *S. mansoni* eDNA, following *in silico* and *in vitro* optimisation of primers. Conventional malacological surveys were conducted in parallel to validate eDNA findings, and statistical analyses examined the influence of environmental factors (turbidity and total dissolved solids (TDS)) on eDNA yield and detection rates.

**Results:** The SYBR Green *S. mansoni* qPCR assay had a practical limit of detection (LOD), defined as amplification in >95% of 33 technical replicates, of 100 copies per reaction. The assay amplified in 82% of reactions with 10 DNA copies and 76% with a single copy. The theoretical LOD, determined via amplification probabilities in RStudio, was 83 copies per reaction, which was also the calculated limit of quantification (LOQ). *Schistosoma mansoni* eDNA was detected in 58.1% (25/43) of filters from Lake Albert and 19.2% (10/52) from Lake Victoria. The Waterra capsule filter yielded the highest eDNA concentrations, while the Sylphium-coupled capsule filter exhibited comparable detection efficiency. Among the DNA purification methods tested, the Sylphium precipitation-based protocol produced significantly higher eDNA yield than column-based kits (DNeasy Blood and Tissue and Zymo Research). Despite significant variation in eDNA recovery across filtration methods, qPCR detection rates were consistent. No significant correlation was observed between turbidity and *S. mansoni* eDNA detectability.

**Conclusion:** Our findings highlight the potential of eDNA as a sensitive, scalable tool for schistosomiasis surveillance. While Waterra filters and Sylphium extraction maximised DNA yield, lower-yield filtration methods still enabled *S. mansoni* detection in high-transmission settings. These findings support the adaptability of eDNA approaches across varying resource contexts. Future work should prioritise protocol standardisation, ecological validation, and development of field-ready diagnostics such as LAMP to enable broader implementation in endemic regions.

## 1. Introduction

Schistosomiasis, a parasitic disease caused by trematodes of the genus *Schistosoma* (WHO, 2016), remains a major public health concern in parts of Latin America, Asia, and particularly Africa, where it leads to significant morbidity and mortality (Aula et al., 2021a; Mawa et al., 2021). The disease has a complex lifecycle that involves freshwater snails as intermediate hosts and humans or other mammals as definitive hosts. In Africa, the two main species affecting humans are *Schistosoma mansoni* and *Schistosoma haematobium* (Aula et al., 2021). According to the World Health Organization (WHO), more than 250 million people require medical treatment for schistosomiasis with over 90% of fatal cases occurring in African countries (WHO, 2016, 2020). In addition, approximately 800 million people in Sub-Saharan Africa remain at risk of infection (Aula et al., 2021).

As the second most prevalent parasitic disease after malaria (Joof et al., 2021; Mawa et al., 2021), there is an urgent need to invest in the eradication of schistosomiasis. To guide this effort, the WHO proposed the 2021-2030 Neglected Tropical Disease (NTD) roadmap, setting ambitious global targets for schistosomiasis elimination. By 2030, the goal is to interrupt transmission and ultimately eliminate the disease in all 78 affected countries (WHO, 2016, 2020). Achieving this task is the complex endeavour influenced by social, economic, political, and cultural factors (Onasanya et al., 2021). Schistosomiasis disproportionately affects impoverished rural communities with limited access to adequate water, sanitation and hygiene (WASH), and healthcare infrastructure (Aula et al., 2021). Addressing these challenges requires a multidimensional, transdisciplinary approach that integrates multiple control strategies. This approach should combine preventive chemotherapy through mass drug administration (MDA) using Praziquantel with snail control and improvements in WASH (Kura et al., 2020; Onasanya et al., 2021; WHO, 2020). Furthermore, promoting behavioural changes within local communities is essential to breaking the disease transmission cycle (Exum et al., 2019).

Effective control of schistosomiasis relies on accurately identifying and targeting high-risk populations, making large-scale infection mapping a critical component of disease surveillance (Trippler et al., 2022). Surveillance tools play an essential role in monitoring infection prevalence in humans, non-human hosts, and intermediate snail hosts, as well as in verifying the elimination of transmission. Traditionally, detection in humans has been performed using the Kato-Katz stool examination for *S. mansoni* and urine filtration for *S. haematobium* (WHO, 2022). However, these methods have limitations, particularly in terms of sensitivity and accuracy at low infection rates, and they are also invasive, time-consuming (Yin et al., 2021) and do not identify transmission sites.

For detecting infections in intermediate snail hosts, conventional techniques include light-induced shedding and crushing methods (Fuss et al., 2020). More recently, molecular diagnostics such as PCR (Hamburger et al., 1998), rapid diagnostic PCR (RD-PCR) (Schols et al., 2019) and loop-mediated isothermal amplification (LAMP) (Gandasegui et al., 2016) have been introduced to detect prepatent infections in the snail intermediate hosts. While these methods are highly sensitive, detecting infections in snails before cercariae are released in into the water, they are labour-intensive. They require malacological sampling, accurate identification of competent snail hosts, and subsequent laboratory-based processing, including cercarial shedding, DNA extraction and amplification. Yet early detection of vector snail species and their associated schistosome parasite species in the environment is critical for schistosomiasis control (Champion et al., 2021; Sengupta et al., 2019). Therefore, there is an urgent need for advanced, on-site pathogen and vector detection tools to identify potential and active transmission sites, provide early warnings, and support timely risk assessments for human health.

Environmental DNA (eDNA) is emerging as a promising tool to address this need. As a non-invasive and scalable approach, eDNA enables highly sensitive detection of target species directly from environmental samples without requiring physical collection of organisms (Jo, 2023). Defined as all genetic material released by organisms into their surroundings (Jo, 2023), eDNA has been widely applied in ecological studies, biomonitoring, and species identification (Bruce et al., 2021). Moreover, its potential for tracking invasive species and pathogens in both terrestrial and aquatic ecosystems makes it a valuable addition to the schistosomiasis surveillance strategies (Bass et al., 2015; Sengupta et al., 2022).

Recent studies have successfully applied eDNA techniques to detect *S. mansoni* in environmental water samples, demonstrating their value in identifying transmission hotspots (Alzaylaee et al., 2020; Sato et al., 2018; Sengupta et al., 2019). However, these efforts have typically relied on a single filtration method or lacked direct comparison of sampling strategies, which may affect DNA yield and detection sensitivity. Furthermore, little attention has been given to optimising qPCR assays for *S. mansoni* detection, particularly with respect to robustness, limit of detection and quantification and specificity across geographically diverse isolates. The choice of filtration technique also remains a key factor influencing eDNA recovery, yet comparative data on filtration performance, cost-efficiency, and field suitability are still limited.

This study aims to address these gaps by: (1) optimising and evaluating the specificity, sensitivity, and robustness of a SYBR Green qPCR assay for *S. mansoni* detection; and (2) comparing the performance and practicality of three different eDNA filtration and purification methods to identify the most cost-effective and sensitive approach for schistosomiasis monitoring. Our research focuses on freshwater bodies in Uganda, Lake Albert and Lake Victoria, where schistosomiasis remains highly endemic, with the broader goal of strengthening eDNA-based molecular surveillance strategies for control programs in similar endemic settings.

## 2. Materials and methods

### 2.1. Ethics statement

Sampling was conducted within the framework of the Uganda Schistosomiasis Multidisciplinary Research Center (U-SMRC), with ethical approval obtained from the Research Ethics Committee of the Uganda Virus Research Institute (GC/127/940), the Uganda National Council for Science and Technology (HS2568ES) and the London School of Hygiene & Tropical Medicine (29203).

### 2.2. Primer design, *in silico* and *in vitro* testing

The existing primer pair of Sato et al. (2018) targeting a 162 bp fragment of the mitochondrial cytochrome oxidase subunit 1 (COI) gene of *S. mansoni* was tested *in silico* using Primer-BLAST analysis against the taxonomic class Trematoda (taxid:6178). We also ran ecoPCR (Ficetola et al., 2010) on 50 trematode mitogenomes sequences obtained from GenBank database (list adapted from Douchet et al., 2022) and 37 COI gene sequences of different *Schistosoma* isolates from different geographical regions of Africa. All the sequences and their accession numbers used for *in silico* testing are provided in the supplementary information file (SI Table 1). Genomic DNA (gDNA) from adult worms of *S. mansoni, S. haematobium, S. bovis, S. rodhaini, S. mattheei* and *S. curassoni,* collected from various geographical regions of Africa and sourced from the Natural History Museum’s Schistosomiasis Collection (SCAN; Emery et al., 2012), was used to assess primer specificity *in vitro* using conventional PCR. In addition, gDNA from a hybrid cross between *S. haematobium* × *S. bovis* and from non-schistosome trematodes known to co-infect the same intermediate snail host (*Biomphalaria*), was obtained from the Royal Museum for Central Africa collection. The non-schistosome trematodes (*Apharyngostrigea pipientis, Euparyphium capitaneum, Drepanocephalus auratus*) were identified as natural co-infections with *S. mansoni* in *Biomphalaria sudanica* during the field sampling.

### 2.3. SYBR Green qPCR assay optimisation and sensitivity testing

To optimise the *S. mansoni* SYBR Green assay, the reactions were performed on CFX96^TM^ C100 Touch^TM^ Thermal Cycler (Bio-Rad), in a 20 µl total volume, with 10 µl SsoAdvanced^TM^ Universal SYBR® Green Supermix (Bio-Rad), 9 µL and 5 µL DNA template. The qPCR cycles were: 30 seconds at 95° C for initial denaturation, 45 cycles of 95° C for 15 seconds and 60° C for 30 seconds each, followed by a melting curve analysis with 15 minutes of temperature rise from 65° C to 95° C with an increment of 0.5° C. The assay’s limit of detection (LOD) and limit of quantification (LOQ) were evaluated using a seven-step 10-fold serial dilution of standards prepared from a 726 bp COI fragment amplified from *S. mansoni* adult worm gDNA. The double-stranded DNA (dsDNA) fragment contained the target region for the Sato qPCR primers used in the study. The 726 bp fragment was amplified by PCR using the Sma-COI-F forward primer (Sato et al., 2018) and the Schiman_COIR reverse primer (Sengupta et al., 2019). Following amplification, the fragment was purified using magnetic bead-based CleanPCR beads (CleanNA) and quantified with the Quant-iT™ PicoGreen™ dsDNA Assay Kit following the manufacturer’s instructions. The resulting concentration was used to calculate the standard DNA copy numbers using the following equation:

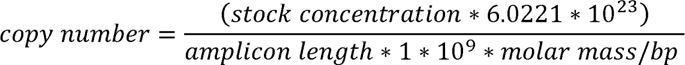

where stock concentration = measured concentration in ng/µL, amplicon length = 726 bp and molar mass of 1 DNA base pair = 650g/mol/bp. The standards, with known copy numbers, were stored at 4°C and used within three days of production to limit any potential effects of DNA degradation. The LOD was assessed both practically and theoretically. Practical LOD was defined as amplification of the lowest standard concentration in >95% of the reactions, while theoretical LOD was the amplification probability for different qPCR replicates for each standard concentration calculated in RStudio (R 4.2.0, with packages “drs”, “ggplot2”) using the script modified from Merkes et al. (2019) published by Klymus et al. (2020). All the qPCR reactions were performed in triplicates.

### 2.4. Field surveys in Uganda

#### 2.4.1. Location of the study

#### 2.4.2. eDNA sampling

Three eDNA filtration methods were used: cellulose nitrate open membrane filters (41 µm and 0.45 µm pore sizes); polyether sulfone Waterra capsule (0.45 µm), polyether sulfone Sylphium single (0.8 µm) and coupled capsules (5 µm and 0.45 µm) (Figures 2b, 2c and 2d). Field preparation followed a modified protocol from Laramie et al. (2015). Upon arrival, a field station was set up (Figure 2a), and reusable equipment were decontaminated using 3.75% bleach, rinsed with lake water, and dried. Boots and waders were also disinfected before and after sampling. eDNA filtration was conducted prior to snail sampling to prevent contamination. Water collection and filtration were performed upstream from the sampling location point. A negative control (field blank) was collected at each site using bottled water to monitor potential contamination. To minimize cross-contamination, different single-use gloves were worn at different sampling sites and changed during filter handling. At each site, 3 L of lake water was collected from five randomly selected microhabitats (total 15 L), ensuring variation in vegetation and shoreline distance. This collection was repeated for both the open membrane and Waterra filtration methods, while for Sylphium filters, water was directly filtered while traversing the microhabitats.

**Figure 1:**
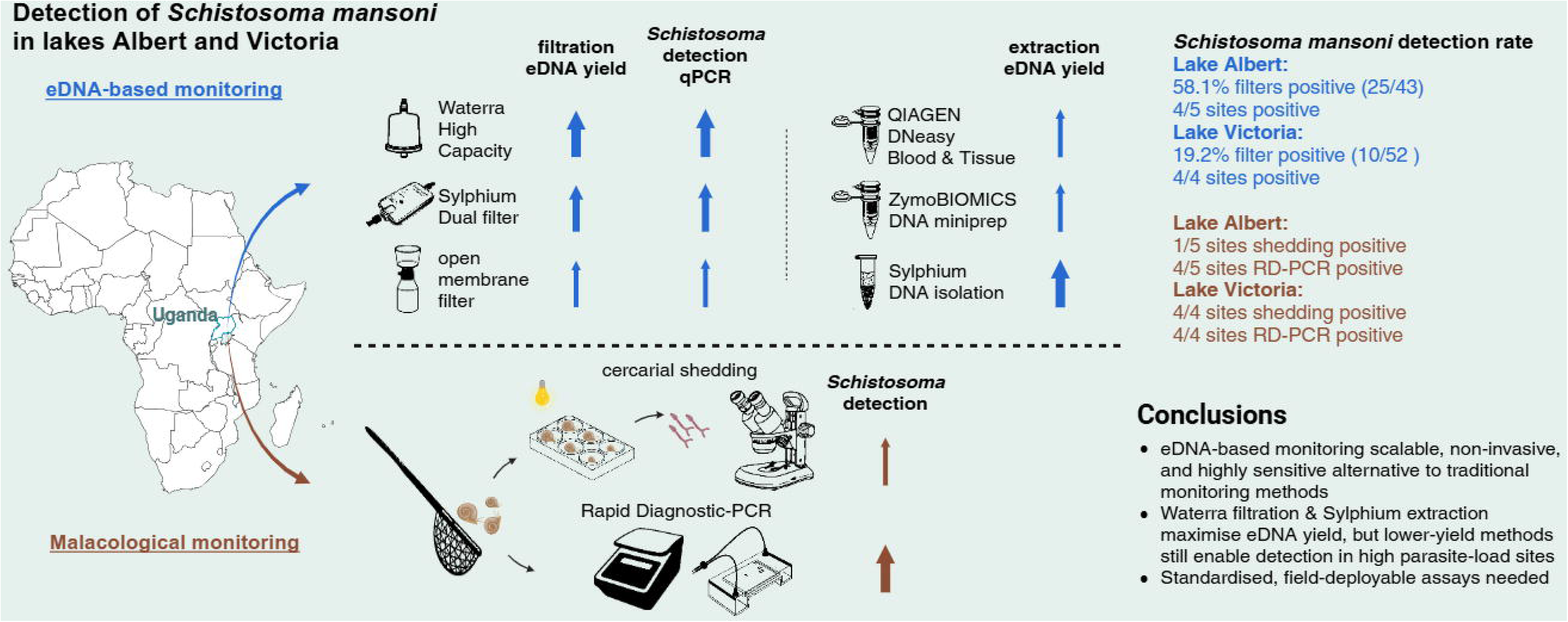
An overview map of the sampling sites in Lake Albert and in Lake Victoria. Along Lake Albert, five sites (Kiina, Sunzu Bay, Sunzu Beach, Songa and Nyalebe) were located at or near the lake shore and two sites (Ngoma and Nyampindu) were inland sites. Along Lake Victoria, five sites (Maruba, Bumeru, Buduma Beach, Busiro Beach and Lugala) were located at or near the lake shore and one site (Kasebere) was an inland site.

**Figure 2:**
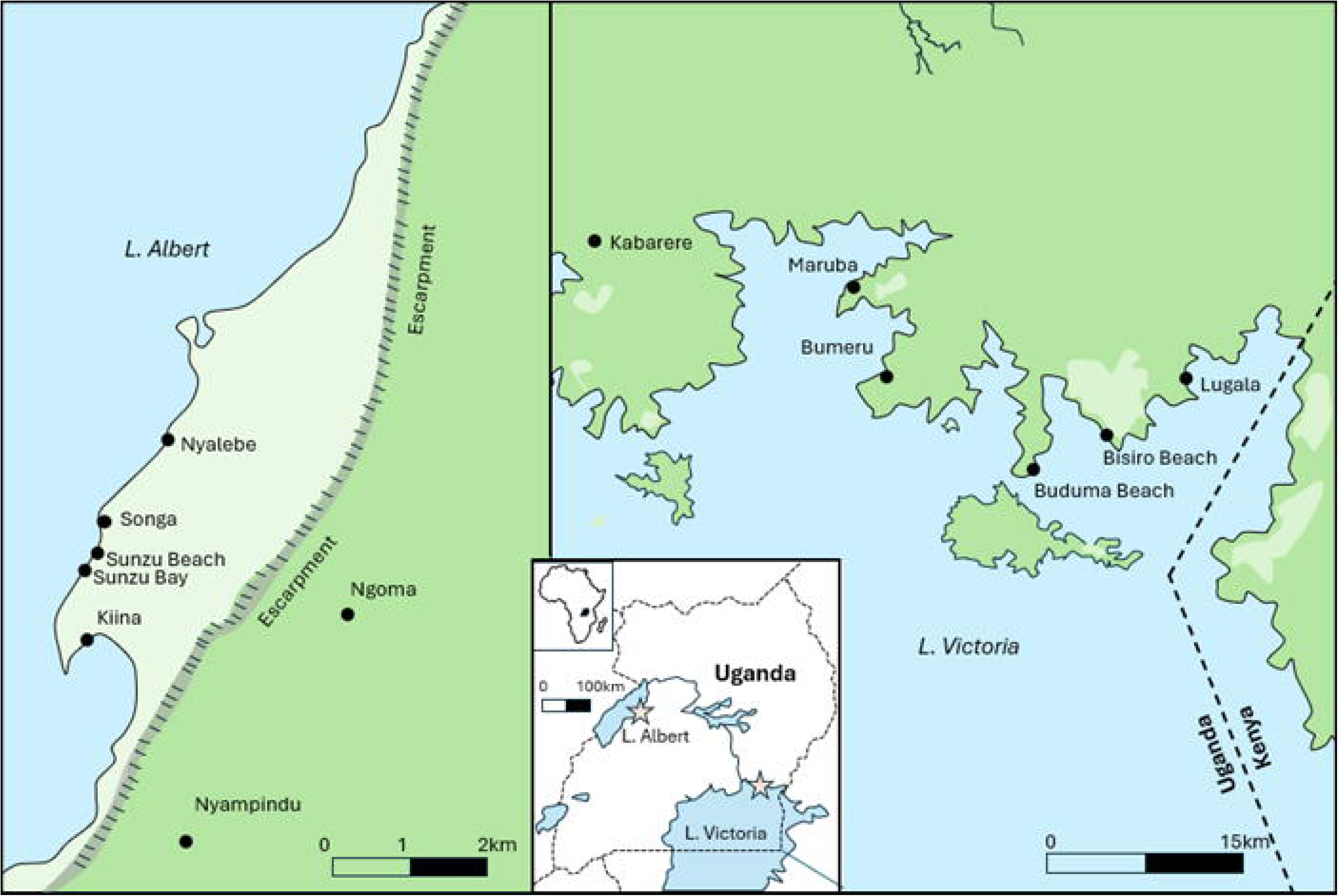
The field setup for the eDNA filtration. a) Decontamination setup with the pink box containing 3.75 % bleach solution. The pink box was also used as a storage box for the equipment to be soaked in bleach before and after sampling, as well as between sampling sessions. The two clear buckets contain water from the sampling site that is used to rinse of the bleach solution after decontamination of the equipment. b) Open membrane eDNA filtration method setup. c) Waterra eDNA filtration setup. d) Sylphium eDNA single filtration setup. The filter capsule is attached to the 60 mL syringe via the silicon valve connector.

##### 2.4.2.1. Open membrane filtration

The open membrane filtration protocol was adapted from Laramie et al. (2015), Protocol #1 with a manual hand-driven vacuum pump. A Thermo Scientific™ Nalgene™ reusable bottle-top filter was used, consisting of a filtration funnel, membrane holder, and a 500 mL plastic receiver bottle to measure filtrate volume. Filtration materials were stored in a closed plastic bucket with 3.75% bleach before and between sites. After decontamination, clean gloves were worn, and the filtration unit was assembled. The membrane filter was attached to the holder and funnel, sealed tightly, and connected to the manual vacuum pump via plastic tubing. Filtration was performed in two steps: first, lake water was passed through the 41 µm membrane, and the filtrate was collected. This filtrate was then filtered through the 0.45 µm membrane. Filtration duration and filtrate volume were recorded. Water was poured slowly into the funnel, maintaining steady vacuum pressure for unidirectional flow. If clogging occurred, the filter was removed with clean forceps and cut in half using decontaminated scissors. One half of the filter was stored in a 2 mL tube with 1.5 mL of DNA/RNA Shield buffer (Zymo Research), while the other half was preserved in a 50 mL Falcon tube with 15-20 grams of silica beads to absorb moisture. Both tubes were sealed with parafilm or duct tape, placed in a labelled Ziplock bag, and stored in a cool box away from light and heat. After decontamination, the same process was repeated for the negative control using bottled water.

##### 2.4.2.2. Waterra filtration

The Waterra eDNA filtration setup included a 20 L jerrycan with a bottom tap, connected to the Waterra filter capsule via plastic tubing. A graded pitcher was placed under the filter outlet to collect and measure the filtrate. Both filtration duration and filtrate volume were recorded. Before use, all equipment was disinfected following the standard decontamination protocol. The jerrycan was disinfected with 3.75% bleach, rinsed three times with lake water, and then filled with water collected from different locations at the sampling site using a clean bucket. Once filled, the jerrycan was placed on a table, and the Waterra filter connected with the flow arrow pointing downward. Gravity-fed filtration was ensured by keeping the jerrycan’s top cap unscrewed to maintain water flow. Once the target volume was filtered or the filter became clogged, the Waterra eDNA filter was detached, and any remaining water inside was carefully removed by shaking the capsule. One end was sealed, and 10 mL of DNA/RNA Shield buffer was added in the opposite direction of the flow arrow. The buffer was mixed by gently shaking the filter. Both ends were then sealed with caps and secured with duct tape to prevent leakage. The filter was placed in a labelled Ziplock bag with the sampling location and date, and then stored in a cool box to protect it from heat and direct sunlight.

##### 2.4.2.3. Sylphium filtration

The Sylphium eDNA filtration kit included a 3.5 mL CTAB preservation buffer (in a 5 mL syringe) containing a xenobiotic internal positive control, along with two luer-lock male red caps (Sylphium Molecular Ecology, Groningen). The filtration method followed the Sylphium Molecular Ecology (2020) protocol with minor modifications. The filter’s valve connector was attached to a 60 mL syringe, and the filter’s inlet submerged in water. Water was drawn into the syringe by pulling the plunger, ensuring minimal air intake to prevent bubble formation that could reduce filtration efficiency. The water was then pushed back out through the filter, and this process was repeated until the filter clogged. Filtration time and volume were recorded. After filtration, residual water in the filter capsule was removed by drawing air through. Once emptied, the filter was disconnected from the syringe, and the outlet was sealed with a cap. To preserve the eDNA sample, 3.5 mL of CTAB buffer was injected from the inlet, and a second cap was secured. The filter was then placed in a labelled Ziplock bag with the sampling location and date and stored appropriately.

#### 2.4.3. Snail sampling, shedding experiments and molecular analysis

Conventional malacology surveillance was also conducted to validate eDNA results. Snail specimens were collected after eDNA filtration (to avoid contamination), with a scooping net (a sieve with 5 x 5 mm mesh mounted on a 2 m metal rod) for 30 minutes at each sampling site by two people following Madsen et al. (2001). Protective gear (gloves and waders) was used to avoid skin-water contact. Both emergent and submerged vegetation were visually inspected for snails. After 30 minutes, the snail specimens were kept alive in empty, clean falcon tubes (50 mL) with some pieces of aquatic vegetation. The snails were further morphologically identified and sorted according to Brown (1994) and the Danish Bilharziasis Laboratory identification keys (Frandsen et al., 1980). The morphological identification was conducted based on shell morphology, shape of soft parts extended out of the shell, i.e., the shape of the foot and the tentacles, head, and presence or absence of an operculum (Brown, 1994; Frandsen et al., 1980).

Each living snail was stored separately in Falcon® 24-well clear flat bottom TC-treated multi-well cell culture plates, filled with bottled water, kept in the dark overnight and exposed to bright artificial light to induce the shedding of cercariae. The snails were examined under the light microscope to detect any cercariae that emerged. When shedding snails were detected, they were stored separately in 97% ethanol with the cercariae in a 15 mL glass tube to keep the shedding snail and the corresponding cercariae together. The tubes were labelled with the location, the date, the snail species, and if the shedding cercariae belong to the genus *Schistosoma* or not based on the first visual inspection. The samples were then packed and ready for transportation for further molecular analysis.

Snail DNA extraction was carried out using the E.Z.N.A.® Mollusc DNA Kit (OMEGA bio-tek, Inc.) following the manufacturer’s protocol. A total of 136 snails from 10 sampling sites were subsequently used for molecular analysis. These snails included different morphotypes of *Biomphalaria* and *Bulinus* across the selected sites from both Lake Albert and Lake Victoria regions. Snail species were identified based on molecular barcoding by amplifying a fragment of the COI gene, followed by PCR product purification and Sanger sequencing, using primers and protocols described by Hammoud et al., 2022). To assess snail infection prevalence, a two-step molecular diagnostic approach was used. First, a general infection RD-PCR was performed to detect trematode infections, including *Schistosoma* spp., in snail tissue DNA. This infection RD-PCR included three markers: (1) an internal control confirming successful PCR amplification, (2) trematodes-specific marker and (3) a *Schistosoma* spp. marker. Second, samples with a positive schistosome signal were then analysed using a multiplex *Schistosoma* RD-PCR to identify the schistosome species up to species level (Schols et al., 2019).

#### 2.4.4. eDNA purification and yield measurement

Three different eDNA purification methods were tested on filtrates from the three eDNA filter types. (1) The Sylphium Environmental DNA Isolation Kit (Sylphium molecular ecology, 2024) was used for Sylphium eDNA single and coupled filter capsules, Waterra filters, and open membrane filters stored in DNA/RNA Shield and dry beads, following the manufacturer’s protocol. Filters stored in dry beads in the field were first placed in 1.5 mL of DNA/RNA Shield solution a few days before purification. For Sylphium filters, 2 mL of CTAB buffer was purified, while 2 mL and 1 mL of DNA/RNA Shield were used for Waterra and open membrane filters, respectively. (2) The DNeasy® Blood & Tissue Kit (QIAGEN) was applied to Waterra filters, where 1 mL of DNA/RNA Shield solution was processed following the manufacturer’s guidelines. (3) The ZymoBIOMICS™ DNA Miniprep Kit (Zymo Research) was used for Waterra and open membrane filters, processing 1 mL of DNA/RNA Shield solution according to the manufacturer’s protocol. For all purification methods, purified DNA was eluted in a final volume of 100 µL, quantified using the Quant-iT™ PicoGreen™ dsDNA Assay Kit following the manufacturer’s instructions and stored at −20°C until further analysis.

### 2.5. Statistical analysis

#### 2.5.1. Data standardisation

To accurately compare the performance of different filters and extraction protocols, the raw DNA yield data and mean Ct values were standardised to account for variations in ratio of volume of the DNA preservative injected to volume extracted. This volume standardisation ensured comparability across filter types and extraction protocols, thus enabling direct comparisons across samples.

Shapiro-Wilk tests were performed to assess the normality of the data distribution. In cases where the assumption of normality was violated, a log-transformation was applied to the data. If normality was not achieved post-transformation, Welch’s ANOVA (Analysis of Variance) or a non-parametric Kruskal-Wallis test was used instead of parametric test. *Post-hoc* comparisons were performed using Tukey’s Honestly Significant Difference (HSD) test for parametric data and Dunn’s test for non-parametric data with Bonferroni adjustments.

The relationship between turbidity (measured in Formazin Nephelometric Units, FNU) and total dissolved solids (TDS, measured in ppm) was assessed to determine whether both environmental variables should be included as covariates in subsequent analyses. Spearman’s rank correlation was used to evaluate the non-parametric relationship between turbidity and TDS.

#### 2.5.2. Comparison of filter performance and DNA purification protocols

To compare the performance of different filter types while accounting for potential environmental variation, a Welch ANCOVA (Analysis of Covariance) was performed. Turbidity and TDS were treated as covariates in the model to assess their influence on DNA yield. The analysis also examined potential interactions between filter type and environmental conditions (turbidity and TDS) through an interaction test. To compare the performance of different DNA purification protocols, a one-way ANOVA was conducted.

#### 2.5.3. Impact of filter type, site and turbidity on *S. mansoni* eDNA detection

A two-way ANOVA was conducted to evaluate the effect of filter type and sampling site on qPCR cycle threshold (Ct) values. This analysis aimed to determine whether different filter types or site conditions influenced the detectability of *S. mansoni* eDNA. Spearman’s rank correlation was used to assess the association between turbidity and eDNA detectability.

All analyses were conducted in R (v4.1.2). Data standardisation and manipulation were performed using the *dplyr* package, while data visualisation (boxplots, scatterplots, histograms, beeswarm and heatmap) was achieved using *ggplot2*. Welch ANOVA and ANCOVA analyses were conducted using the *car* package, and *post-hoc* tests were performed using emmeans and *FSA* packages.

## 3. RESULTS

### 3.1. Assay specificity and sensitivity

The *in silico* test showed 100% specificity of primers for conspecific mitolineages. The conventional PCR results presented a high specificity to *S. mansoni* without cross-amplifying other trematode DNA, including other *Schistosoma* species (Figure 3). The observed or practical LOD was 100 copies per reaction. The assay amplified in 82% of the reactions with ten copies of DNA and in 76% with one copy of target DNA (Table 2). The theoretical LOD and calculated LOQ was 83 copies per reaction (Figure 4).

**Figure 3:**
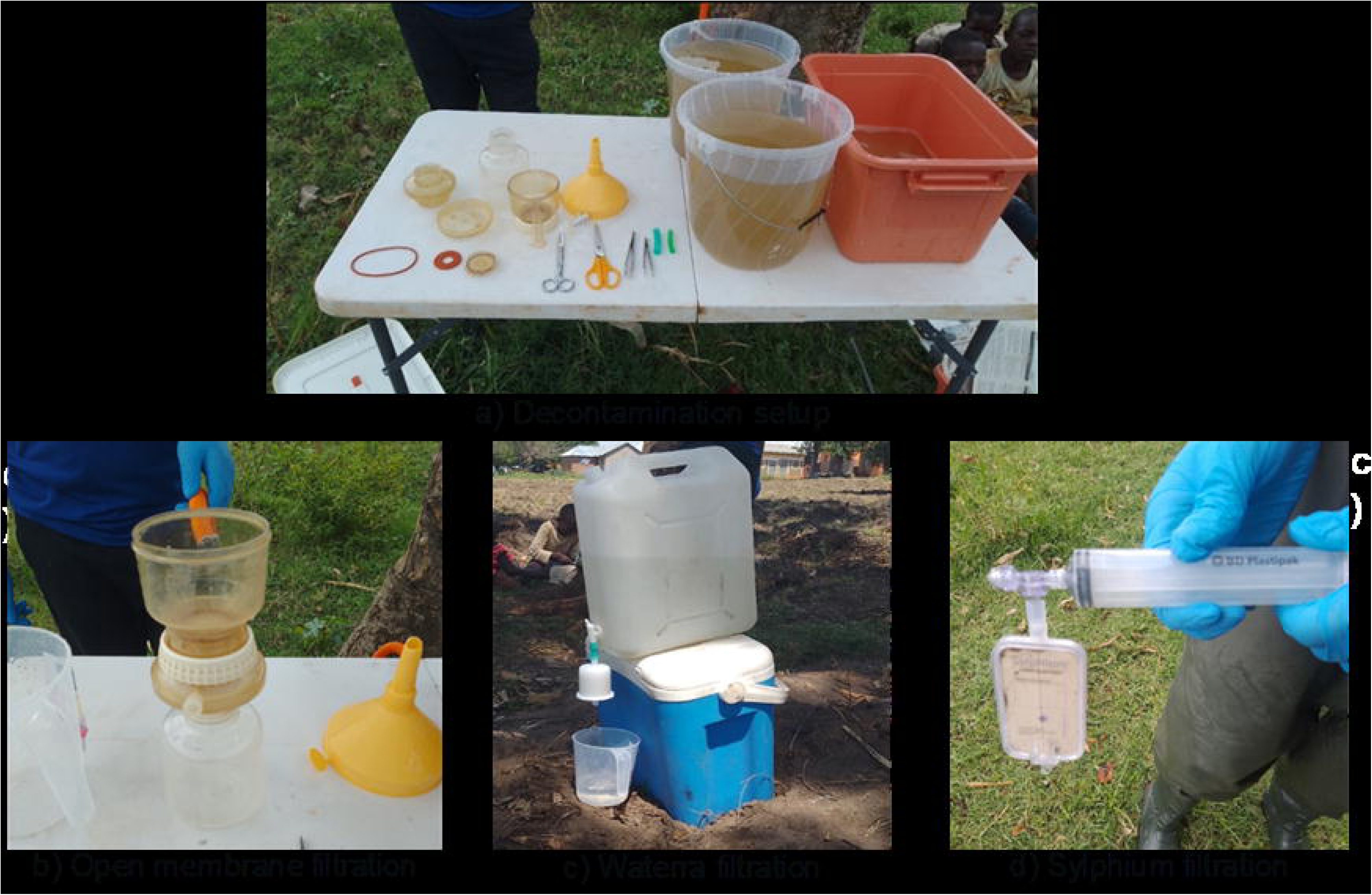
Results of the *in vitro* specificity test for *S. mansoni* primer pair. (A) gDNA of *Schistosoma* species isolates: lanes 1, 2, 3 and 7 contain the gDNA of *S. mansoni* isolates from Cameroon, Egypt, Kenya and Senegal, respectively; lanes 4, 5, 6 and 8 contain *S. haematobium* isolates from Liberia, Zanzibar, Guinea Bissau and Senegal, respectively; lane 9 is *S. bovis* isolate from Kenya, lane is 10 *S. rodhaini* isolate from Burundi, lane 11 is *S. mattheei* isolate from Zambia and lane 12 is *S. curassoni* isolate from Senegal. (B) gDNA of infected and non-infected *Biomphalaria* and *Bulinus* snails isolated from Uganda: lane 1,2,3 are *Bi. Sudanica snails infected with Drepanocephalus auratus, Apharyngostrigea pipientis* and *Euparyphium capitaneum respectively*, 4, 5, 6 are *Bu. truncatus*, 7-11 are *Bi. sudanica* (7,8,10,11 infected with *S. mansoni* and 9 non-infected), and lane 12 contains a negative PCR control (UltraPure^TM^ DNase/RNase Free distilled water). L: DNA ladder (100 bp each).

**Figure 4:**
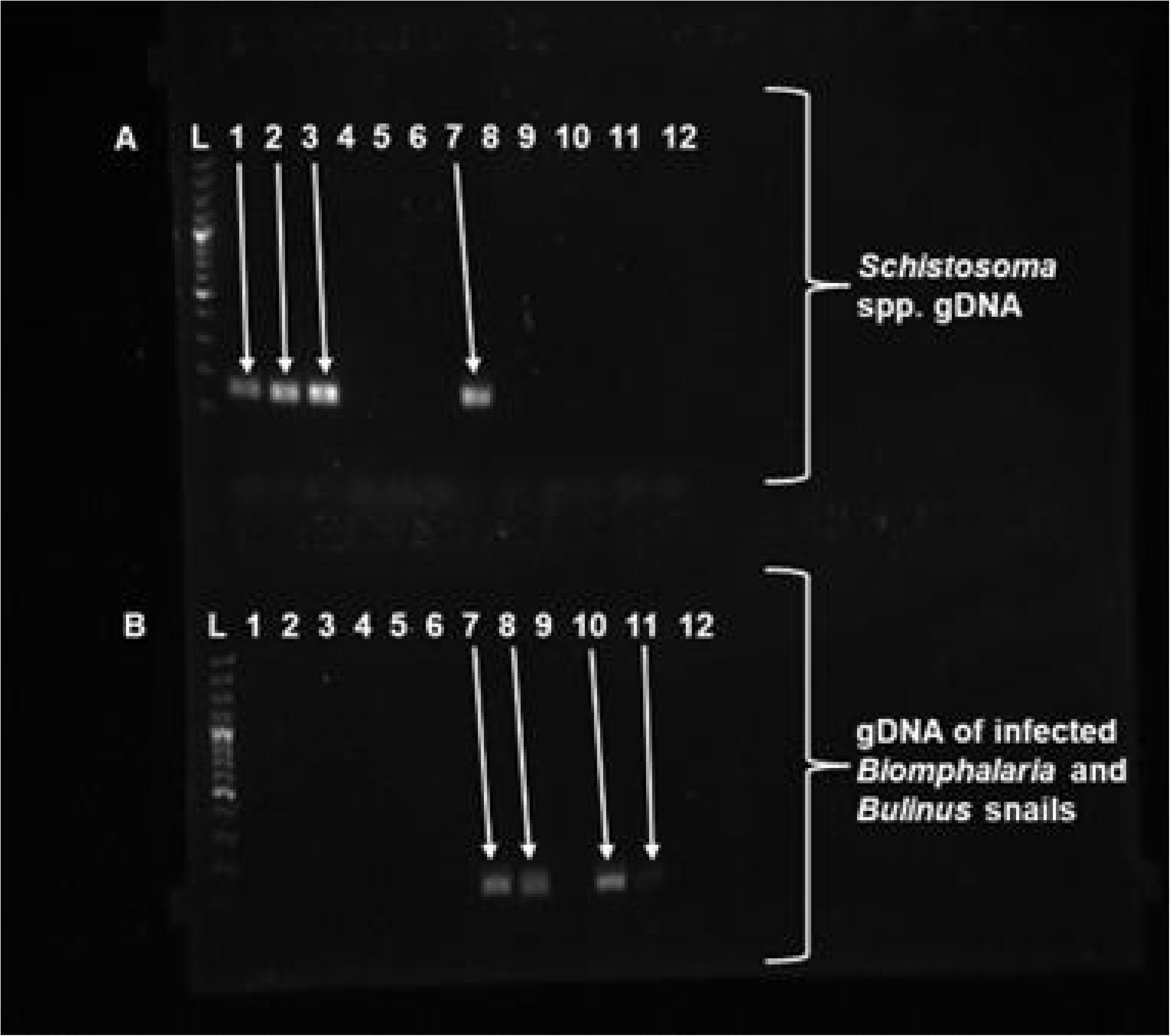
(a) The Calibration curve plot with Ct values on the y-axis and standard concentration on the x-axis. The black points are the middle 2 quartiles of standards with ≥50% detection and are included in the linear regression calculations, while the blue pluses (+) are not included in the linear regression calculations as they fall outside the middle 2 quartiles and/or with <50% detection. The mean R^2 of the assay is 0.979. The resolved limit of detection (LOD) and limit of quantification (LOQ) are indicated by the vertical red and black dotted lines, respectively, and show a value of 83 copies/reaction. (b) The theoretical LOD plot showing the probability of successful amplification at different qPCR replicates for each standard concentration. Each colour represents different LOD for different numbers of qPCR replicates. (c) The LOQ plot for the assay. The y-axis represents the coefficient of variation (CV) for Ct values, while the x-axis shows standard concentrations in copies per reaction. The blue line represents the LOQ model, and the points indicate the CVs for each standard concentration. The vertical red line marks the LOD for reference. The grey rectangle represents the calculated LOQ, with its right boundary indicating the value (83 copies/reaction), determined where it intersects the calibration curve (LOQ is similar to the LOD in this study). The plots are produced in RStudio, code modified from Merkes et al. (2019).

**Figure 5:**
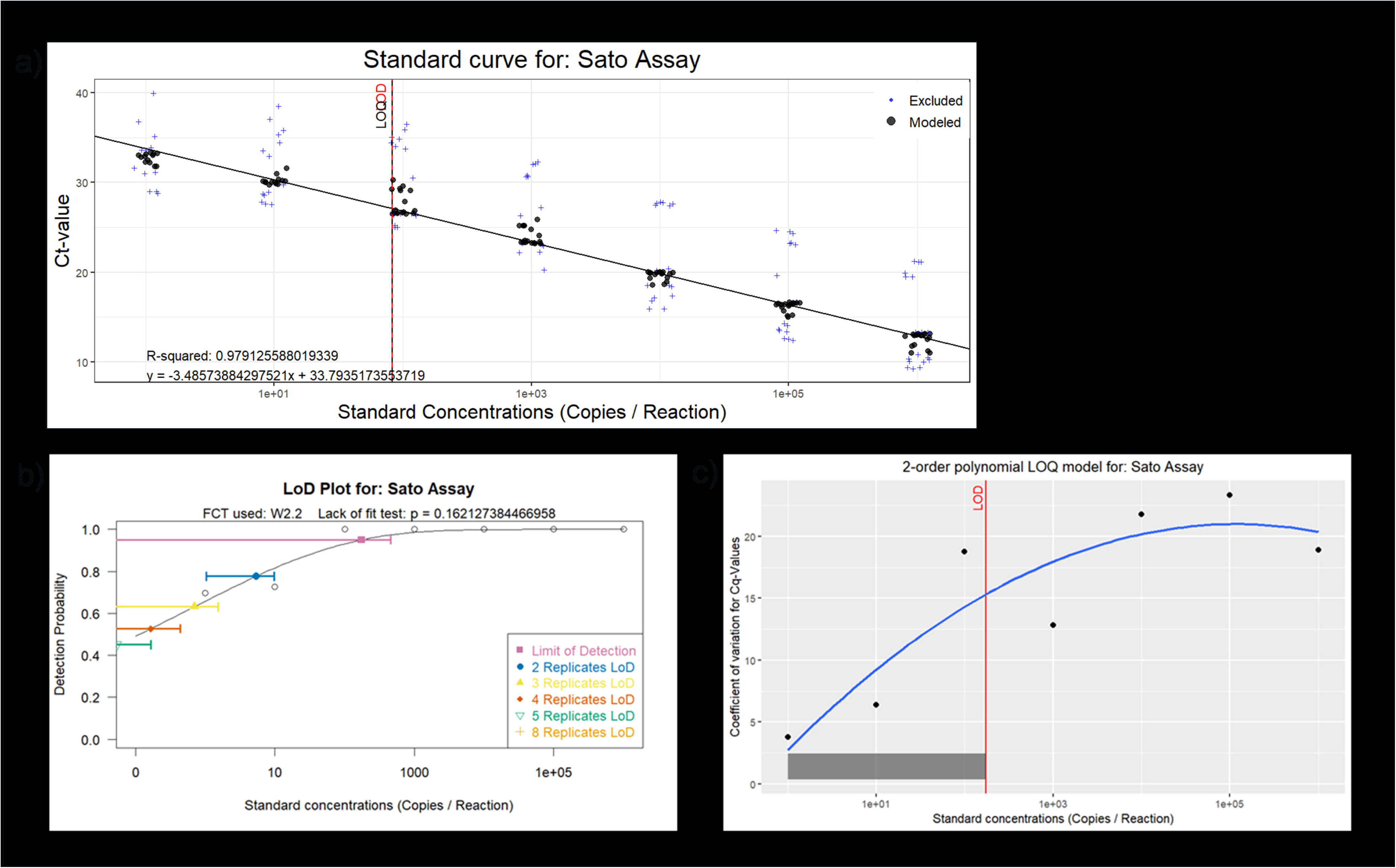
Filter type performance based on a) total eDNA yield captured and b) volume of water filtered. The black line represents the mean and the red line the median volume.

**Table 1:**
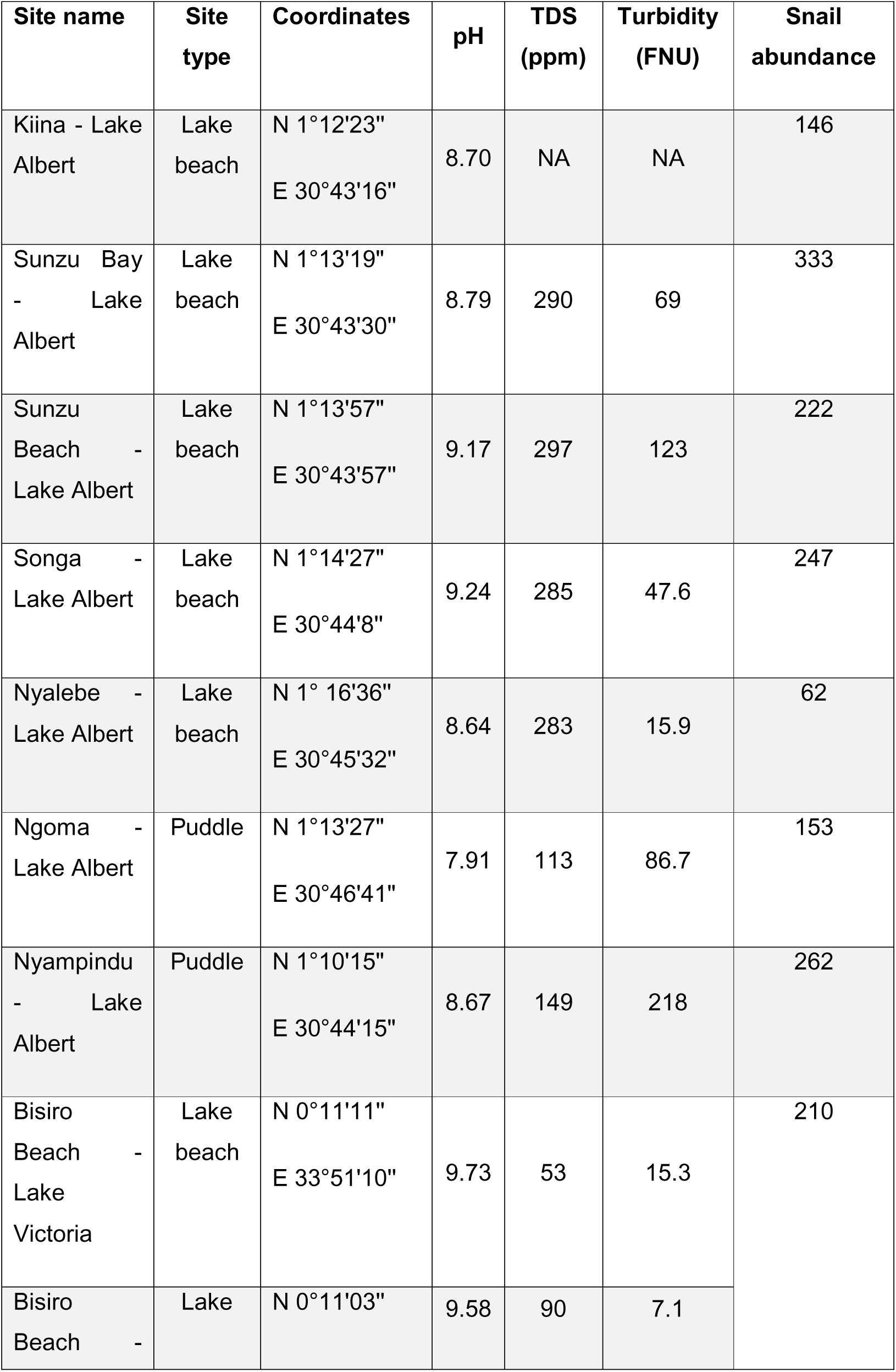
Sampling sites, geographical coordinates, biotic and abiotic parameters.

**Table 2:**
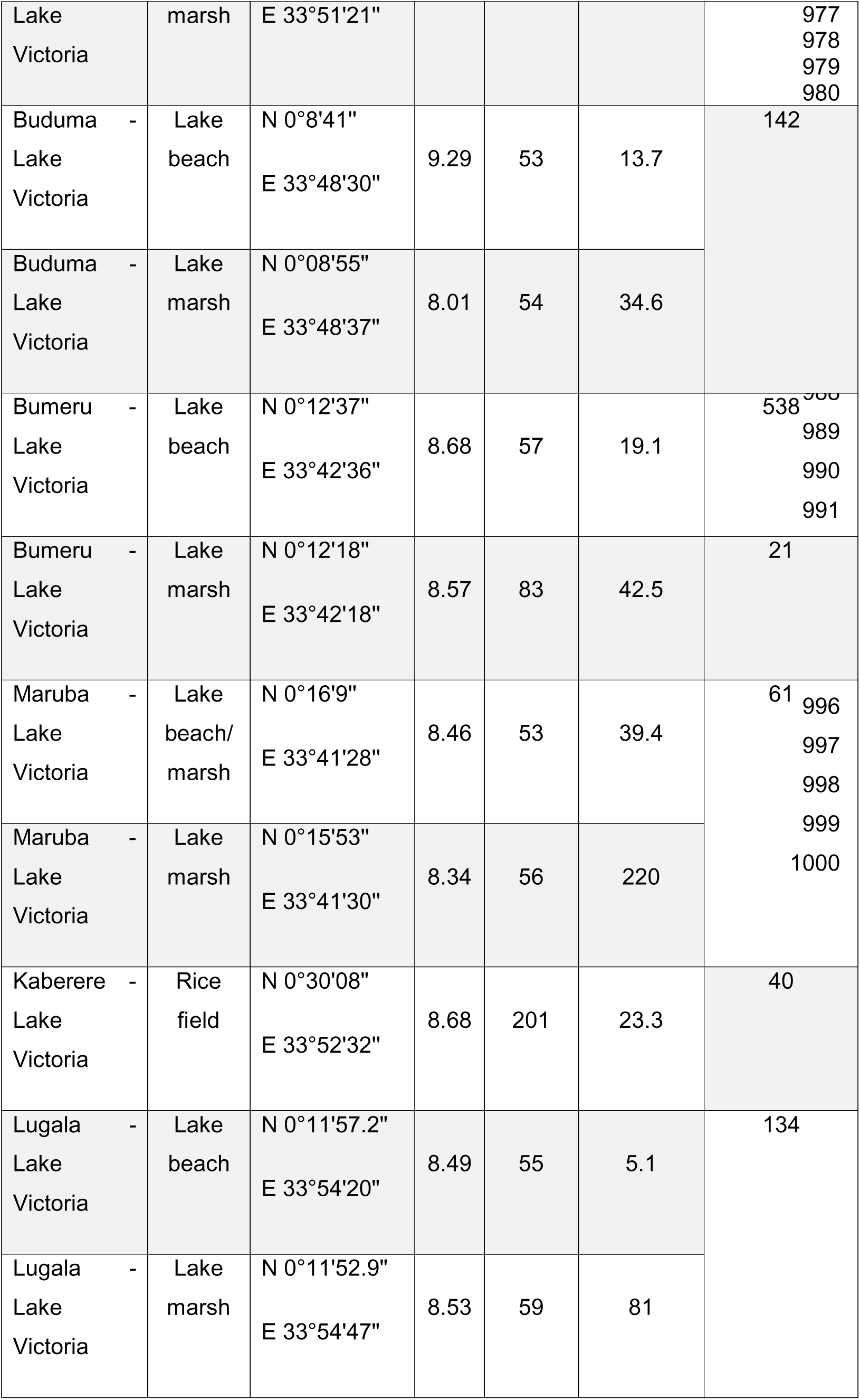
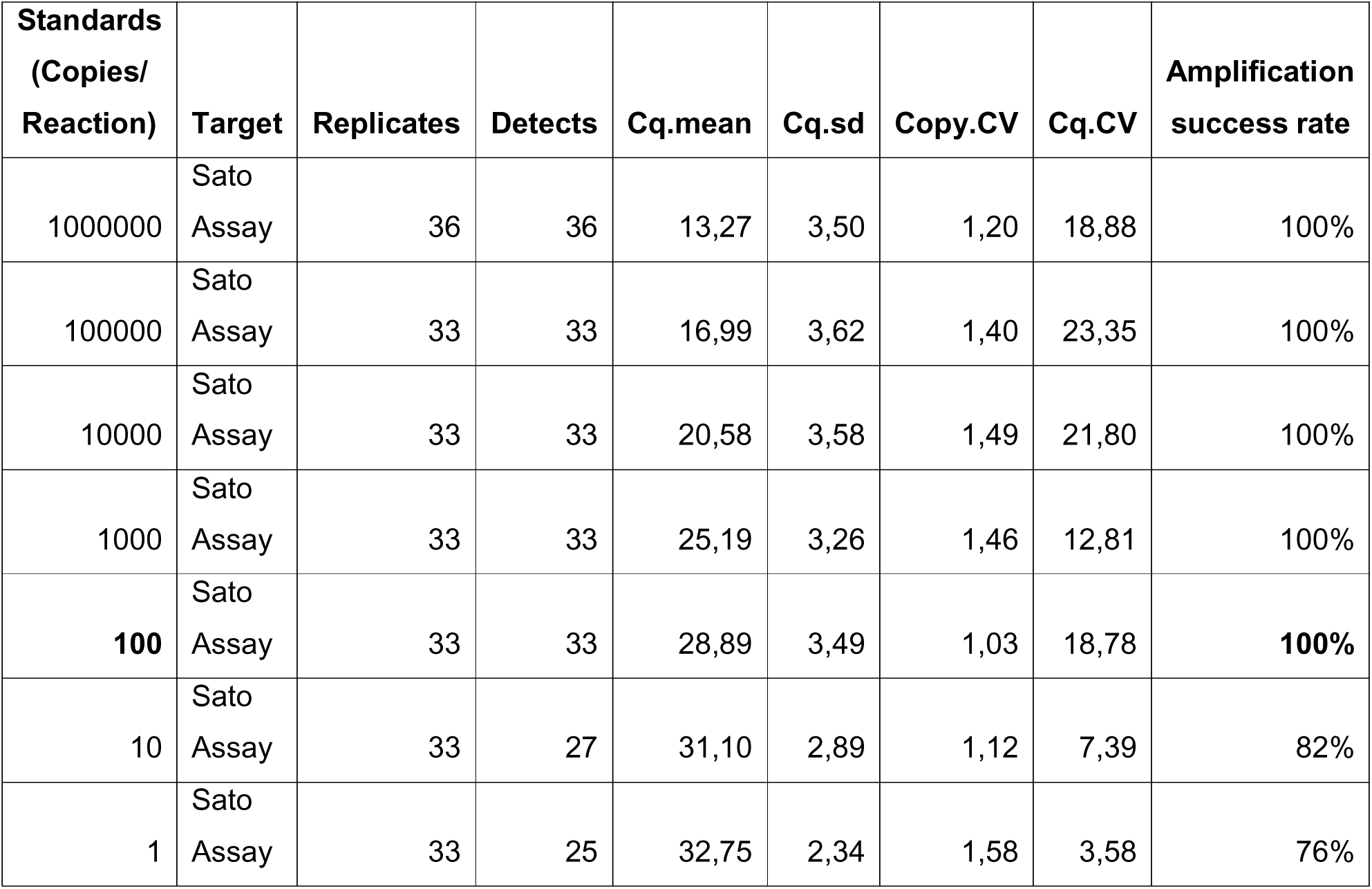
Amplification success rate and mean Ct values for individual standard concentrations.

### 3.2. eDNA field samples qPCR amplification

A sample was confirmed positive for *S. mansoni* parasite eDNA when the melt peak obtained for one or more replicates was 77°C ± 0.5°C, defined by the melting curve profile of the positive controls. *Schistosoma mansoni* eDNA was detected in 25/43 (58.1%) filters (including the field filtration and laboratory extraction negative controls) analysed from Lake Albert and 10/52 (19.2%) from Lake Victoria. Comparing eDNA results with conventional snail surveys, eDNA analysis showed partial agreement with conventional snail survey methods in Lake Albert. Specifically, eDNA results were consistent with RD-PCR in 3 out of 5 sites and with shedding experiments in 1 out of 5 sites (Table 3). Sites that tested positive for *S. mansoni* eDNA but negative in both shedding experiments and RD-PCR suggest that the detected eDNA likely originated from parasite eggs and miracidia present in the environment rather than from cercariae. Conversely, sites that were negative for eDNA and shedding experiments but positive for RD-PCR indicate that snails at these locations may have been in the early stages of infection (prepatent infection), where they had not yet started shedding cercariae. In contrast, all sites surveyed in Lake Victoria showed complete agreement across eDNA, shedding experiments and RD-PCR. It is worth noting that 5/37 field filtration negative controls showed a positive signal with measurable Ct values. However, they displayed unspecific melting curves compared to the *S. mansoni* positive control melt peak. None of the six extraction blanks and 15 qPCR negative controls displayed measurable Ct values, indicating that the positive field filtration negative controls were likely random amplification and not contamination with *S. mansoni*. All the positive eDNA samples were sequenced as an additional validation step, and the results confirmed that the qPCR amplified eDNA fragments belonged to *S. mansoni*, with an identity ranging from 95.2% to 100% to the matching sequences in the NCBI (accession numbers: MK172832, MT994261, MZ660623, MZ656796, MK095609, MN603504, MF919423, MN593408).

**Table 3:** Comparison of environmental DNA (eDNA) results with cercariae shedding experiments in the field and rapid-diagnostic polymerase chain reaction (RD-PCR).

| **Lake Albert** |  |  |  |  |  |
| --- | --- | --- | --- | --- | --- |
| **Site name** | Kiina | Sunzu<br>Bay | Sunzu<br>Beach | Songa | Ngoma |
| **eDNA** | + | + | + | + | - |
| **Shedding**<br>**experiments** | - | - | - | + | - |
| **RD-PCR** | - | + | + | + | + |
| **Lake Victoria** |  |  |  |  |  |
| **Site name** | Bisiro<br>Beach | Bumeru | Maruba | Kaberere |  |
| **eDNA** | + | + | + | + |  |
| **Shedding**<br>**experiments** | + | + | + | + |  |
| **RD-PCR** | + | + | + | + |  |

**Table 4:**
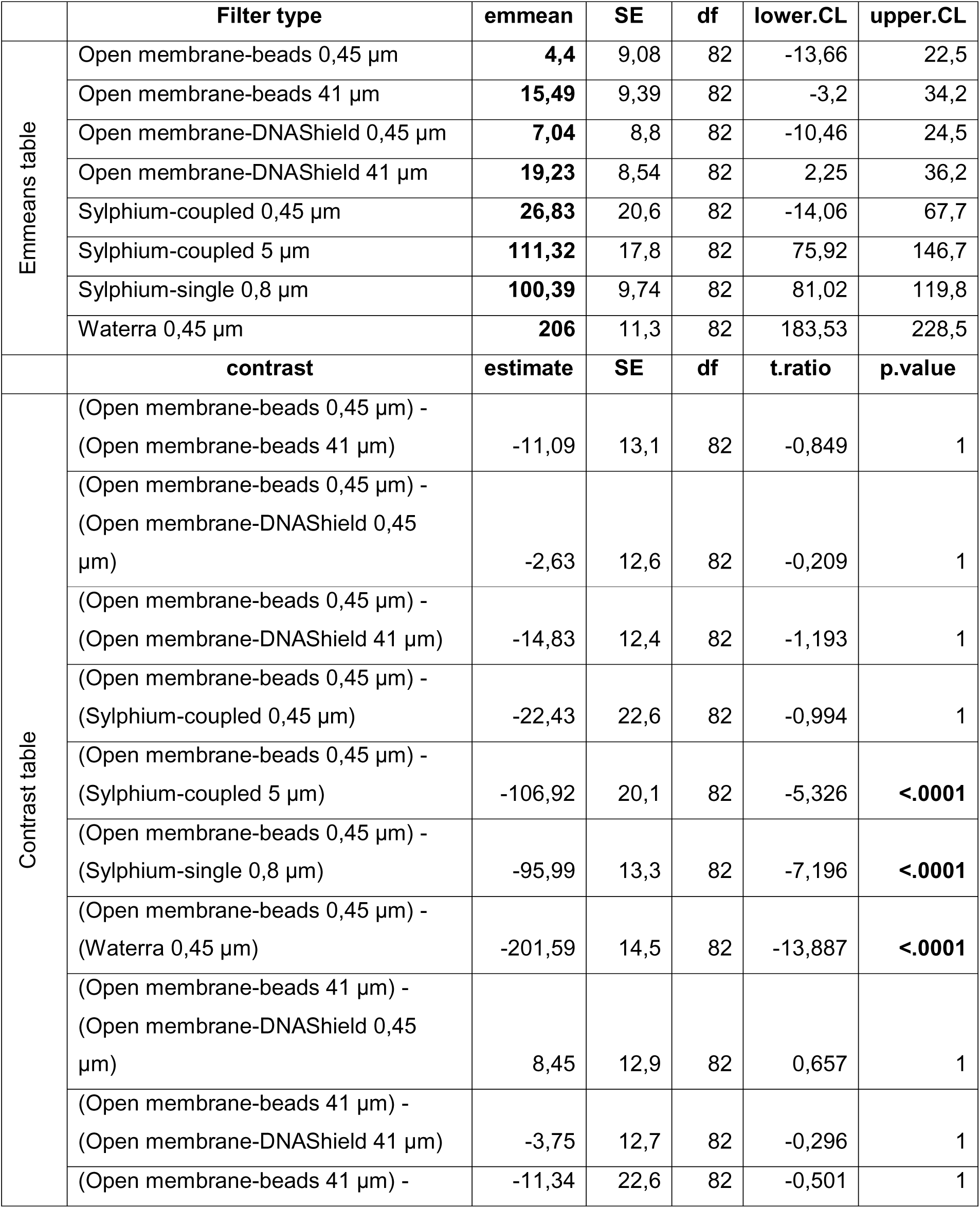
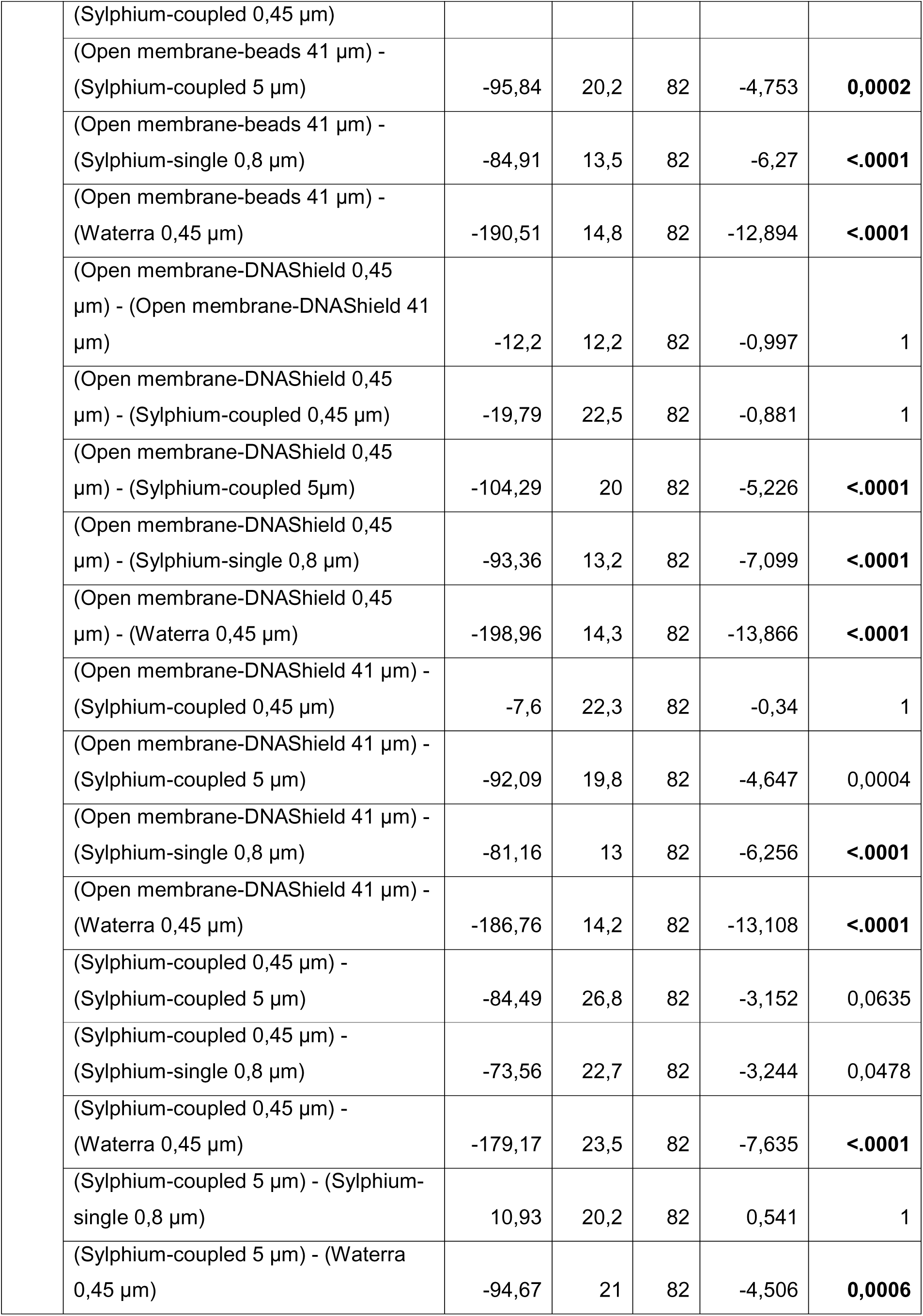
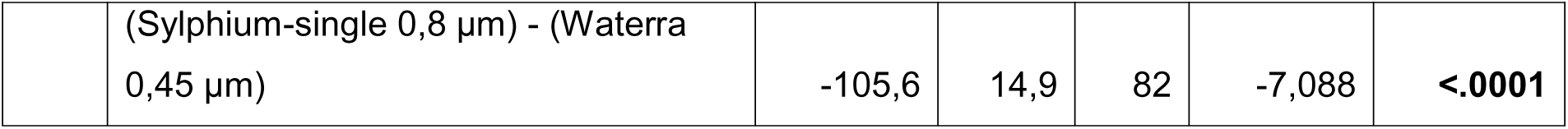
Estimated marginal means (emmeans) and pairwise comparisons of eDNA yield across different filter types. The emmeans table shows the estimated means (emmean) for each filter type along with standard error (SE), degrees of freedom (df), and 95% confidence intervals (lower.CL and upper.CL). The ‘contrast’ table presents pairwise comparisons between filter types, displaying the estimated difference (estimate), standard error, t-ratio, and adjusted p-values using the Bonferroni method. Significant differences (p < 0.05) highlight filters with higher DNA yields.

### 3.3. Correlation between turbidity and TDS

Spearman’s rank correlation analysis revealed a moderate positive correlation between turbidity and TDS (ρ = 0.463, p-value = 1.065e-08). This indicates that as turbidity increases, TDS tends to increase as well, though the relationship was not particularly strong. The significant p-value (p = 1.065e-08) suggests that the correlation is unlikely to have occurred by random chance. The moderate correlation (ρ = 0.463) implies that while turbidity and TDS reflect related environmental factors, they capture distinct aspects of water quality. Therefore, both turbidity and TDS were retained in downstream analyses to avoid overlooking independent contributions to DNA yield variability.

### 3.4. Comparison of filtration techniques across sampling sites (filter performance)

A total of 100 filters from 13 sampling sites were used for the Welch ANCOVA. The results indicated that filter type had a significant effect on eDNA yield (p < 2.2e-16), while neither turbidity (p = 0.174) nor TDS (p = 0.187) demonstrated a significant influence on eDNA yield. This suggests that while filter type is a key determinant of total eDNA yield, the variability in turbidity and TDS does not strongly influence eDNA capture efficiency across the filter types tested. Consequently, an interaction test was conducted to evaluate whether filter performance varied with turbidity and TDS. The results showed no significant interaction between filter type and turbidity (p = 0.503) or TDS (p = 0.078). This indicates that neither turbidity nor TDS significantly altered the performance of the filters in relation to eDNA yield. The lack of interaction suggests that the influence of turbidity and TDS on DNA yield is consistent across filter types.

A *post-hoc* pairwise comparison of total eDNA yield across different filter types was conducted using estimated marginal means (emmeans) and the Bonferroni adjustment for multiple filter types comparison (Table 1). The results indicate that Waterra 0.45 µm had the highest eDNA yield (emmean = 206 ng), significantly outperforming Sylphium and open membrane filters (p < 0.05). Sylphium-coupled 5 µm and Sylphium-single 0.8 µm rank second and third with emmean of 111.32 and 100.39 ng of DNA yield, respectively. These findings highlight the superior performance of Waterra over Sylphium and open membrane filters in eDNA recovery from water samples. The resulting high yield in Waterra filters might be attributed to the filter’s capacity to filter larger water volumes (Figure 2). In contrast, no significant differences were observed in eDNA yield between the open membrane filters with 41 µm and 0.45 µm pore size (p = 1); or between the open membrane filters preserved in Zymo DNA/RNA shield and those preserved in dry beads (p = 1). These results indicate comparable performance in eDNA recovery across the open membrane filters, regardless of pore size or preservation method.

### 3.5. Comparison of eDNA purification methods for Waterra filters

A total of 29 Waterra filters from nine sampling sites were analysed to evaluate the effectiveness of three eDNA purification protocols: DNeasy Blood and Tissue (seven samples), Sylphium extraction (11 samples), and Zymo Research (11 samples), based on eDNA yield. Results from Shapiro-Wilk test indicates no significant deviation of the data from normality for all three methods (p > 0.05) (Table 5). However, the DNeasy Blood and Tissue and Zymo Research yield data showed potential skewness (SI Figure 3). To ensure consistency across all groups, the data was log transformed and Kruskal-Wallis test chosen, followed by a *post-hoc* Dunn’s test to evaluate pairwise differences in eDNA yield between the three purification protocols. The results in Table 6 show significant differences in eDNA yield between DNeasy Blood and Tissue and Sylphium extraction (*Z* = −4.08, p.adj = 0.00013), with Sylphium extraction protocol yielding significantly more eDNA. Additionally, Sylphium extraction showed significantly higher eDNA yield compared to Zymo Research (*Z* = 2.54, p.adj = 0.034). However, no significant difference was detected between DNeasy Blood and Tissue and Zymo Research (*Z* = −1.76, p.adj = 0.236). Figure 6 visually illustrates the differences in eDNA yield across three purification methods after log transformation.

**Figure 6:**
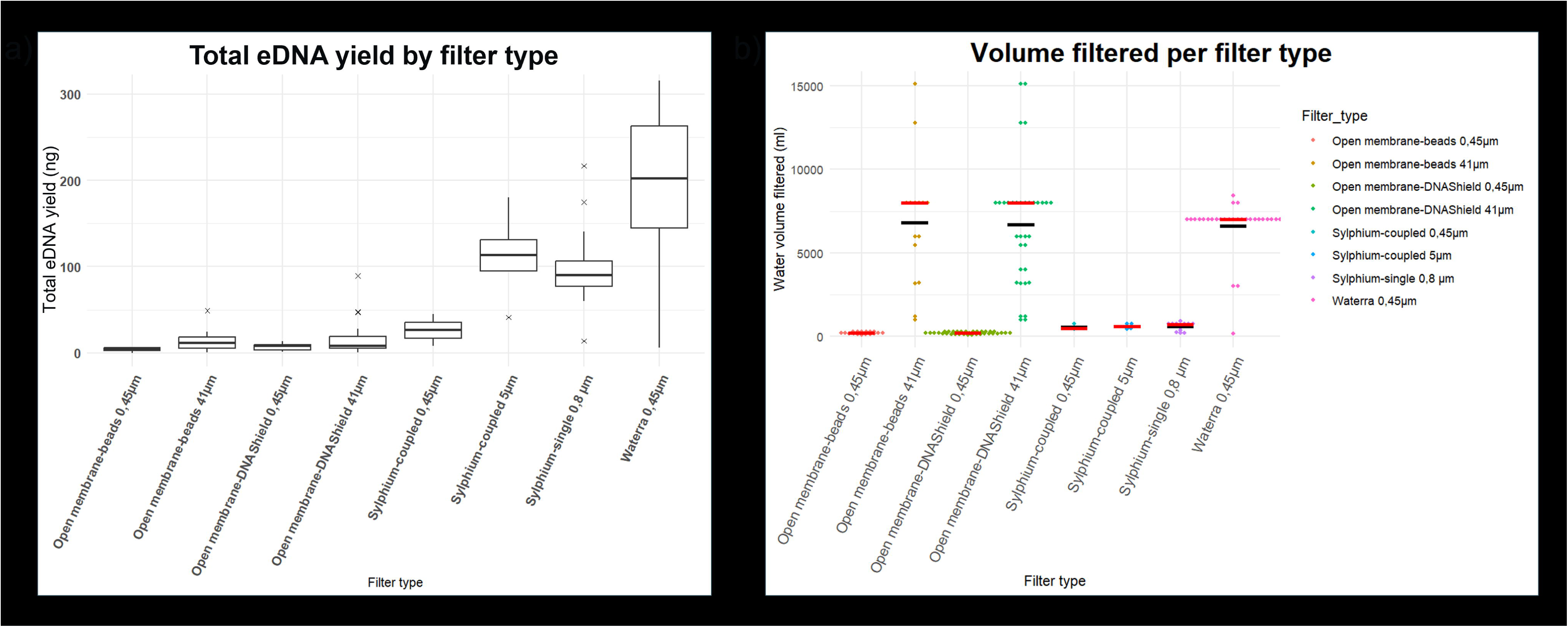
Comparison of eDNA yield across three DNA purification methods: DNeasy Blood and Tissue, Sylphium extraction, and Zymo Research, on Waterra filters from different sampling sites. The y-axis represents the log-transformed eDNA yield (ng/L). Sylphium extraction protocol demonstrates the highest median yield with a wider interquartile range, while DNeasy Blood and Tissue shows the lowest eDNA yield. The box represents the interquartile range (IQR), the horizontal line within the box is the median, and the whiskers indicate variability outside the upper and lower quartiles.

**Table 5:** Shapiro-Wilk normality test results. Shapiro_p is the p-value and n is the sample number.

| DNA purification method | n | shapiro_p |
| --- | --- | --- |
| DNeasy Blood and Tissue | 7 | 0,376 |
| Sylphium extraction | 11 | 0,969 |
| Zymo Reasearch | 11 | 0,802 |

**Table 6:** Dunn *post-hoc* test results comparing eDNA yield across three purification methods: DNeasy Blood and Tissue, Sylphium extraction, and Zymo Research. The results are adjusted for multiple comparisons using the Bonferroni correction to control for Type I error.

| Comparison | Z | P.unadj | P.adj |
| --- | --- | --- | --- |
| DNeasy Blood and Tissue - Sylphium extraction | -4,08497 | 4,41E-05 | 0,000132 |
| DNeasy Blood and Tissue - Zymo Research | -1,75849 | 0,078664 | 0,235993 |
| Sylphium extraction - Zymo Research | 2,536925 | 0,011183 | 0,033549 |

### 3.6. Quantitative analysis by Ct values to assess *S. mansoni* eDNA detectability across sites and filter types

Figure 7 illustrates the variation in Ct values across sampling sites and filter types for *S. mansoni* eDNA detection. Kiina, LA Songa and Sunzu Beach, all in Lake Albert, and Bumeru A and B in Lake Victoria had lower Ct values, indicating higher concentration of *S. mansoni* eDNA detectable by qPCR, likely suggesting higher parasite presence at these locations. In contrast, LV Bisiro A and B, LV Kaberere and LV Maruba A, all in Lake Victoria, and Sunzu Bay in Lake Albert showed higher Ct values, reflecting lower parasite eDNA recovery. Similarly, when comparing filter types, Sylphium and Waterra filters exhibited lower Ct values, suggesting superior detection efficiency compared to open membrane filters. The relatively high Ct values observed with the open membrane filters may indicate reduced *S. mansoni* eDNA filtration efficacy.

**Figure 7:**
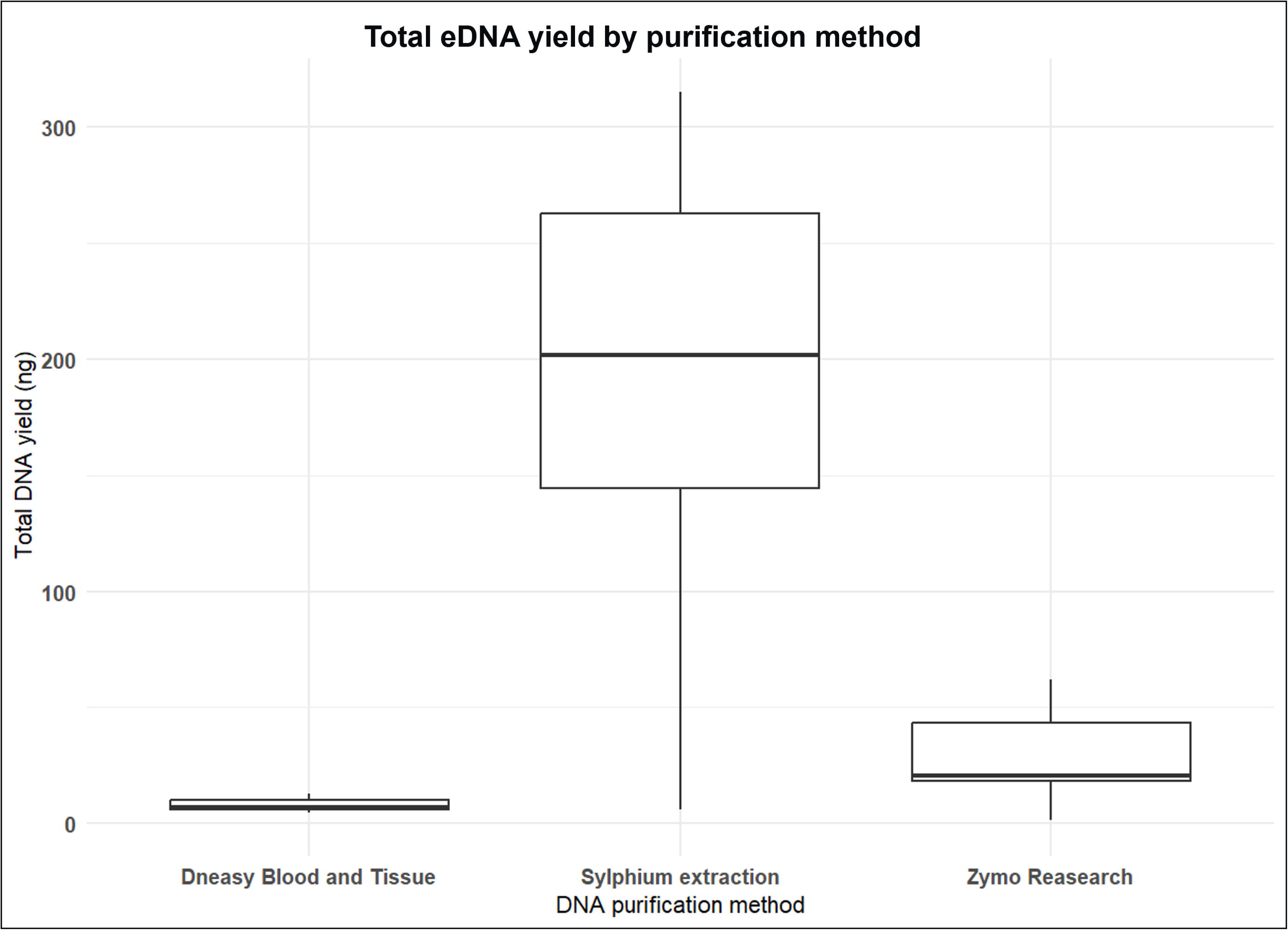
(a) Variation in cycle threshold (Ct) values across sampling sites, indicating potential differences in S. mansoni presence. Sites such as Kiina, Songa, Sunzu Beach (LA) and Bumeru A and B (LV) exhibit lower Ct values, suggesting higher parasite eDNA concentrations, while Sunzu Bay (LA), Bisiro A and B, Kaberere and Maruba A (LV) show higher Ct values, indicating lower eDNA recovery. (b) Distribution of Ct values for S. mansoni detection across different filter types. Sylphium and Waterra filters exhibit lower Ct values, indicating higher detection efficiency compared to open membrane filters.

However, a two-way ANOVA (Table 7) revealed no statistically significant differences in mean Ct values across filter types (*F*(6, 4) = 0.511, p = 0.780) or sampling sites (*F*(8, 4) = 0.505, *p* = 0.809) among samples positive for *S. mansoni* eDNA. This indicates that, despite apparent descriptive trends, the variability in Ct values was not significant when considering filter type or site alone. Nevertheless, whenever there was disagreement between filters, Waterra had higher detection rates (5/6 positive filters), followed by Sylphium (with an overall 9/17 positivity rate, among which 6/10 single filters were positive), with Open Membrane showing the highest rate of false negatives (with an overall 5/19 positivity rate) (Figure 8).

**Figure 8:**
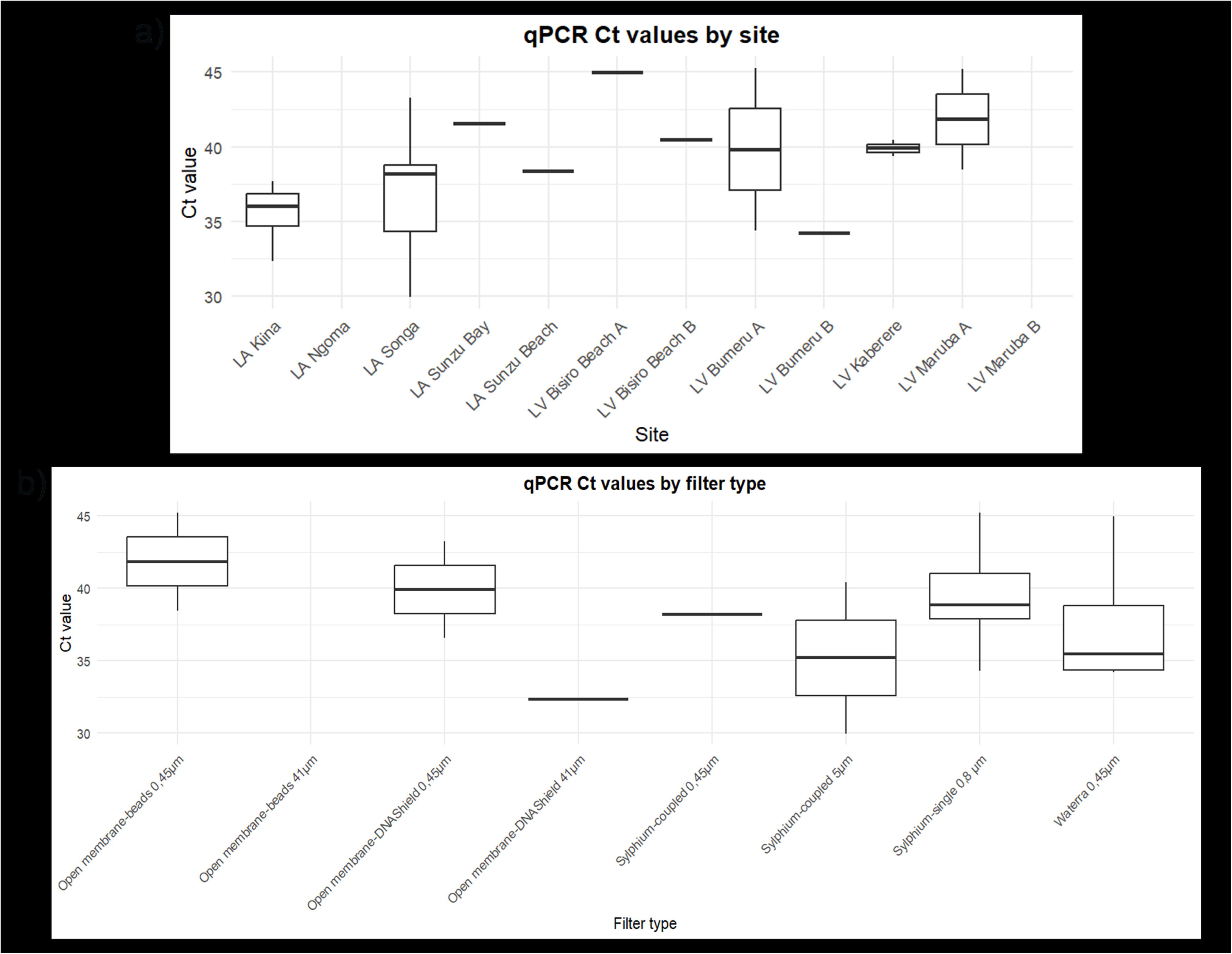
Detection rates for S. mansoni per site, calculated as the proportion of filters testing positive by qPCR.

**Table 7:** A summary of two-way ANOVA results evaluating the effects of filter type and sampling site on mean Ct values for samples positive for *S. mansoni* eDNA.

| Source | Df | Sum Sq | Mean Sq | F value | p(>F) |
| --- | --- | --- | --- | --- | --- |
| Filter type | 6 | 95.85 | 15.97 | 0.511 | 0.780 |
| Site | 8 | 126.37 | 15.80 | 0.505 | 0.809 |
| Residues | 4 | 125.13 | 31.28 |  |  |

### 3.7. Site-level transmission analysis to rank sites based on detection frequency and intensity

Nine out of the twelve sampling sites tested positive for *S. mansoni* eDNA through qPCR analysis. The sites Kiina (LA) and Bumeru A (LV) exhibited the highest detection rates (100%), suggesting active transmission, as all collected filters tested positive for *S. mansoni*. In contrast, Ngoma (LA), and Bisiro Beach B and Maruba B (LV) showed no detectable *S. mansoni* eDNA (0% detection rate), indicating either the absence of transmission or eDNA concentrations below the detection threshold (Figure 8).

### 3.8. Relationship between turbidity and *S. mansoni* eDNA detectability

A weak negative correlation was observed between water turbidity and *S. mansoni* eDNA detectability (Spearman’s rho = −0.359, p = 0.189). This suggests a slight trend in which higher turbidity was associated with lower Ct values, potentially indicating increased eDNA detection. However, the relationship was not statistically significant (p > 0.05), implying that turbidity is not a strong or consistent predictor of *S. mansoni* eDNA detectability in the tested samples.

### 3.9. Analysis of sampling efforts and costs

The time spent in the field and equipment costs were compared among the three eDNA filtration techniques and the conventional malacology method. Although conventional malacological survey was significantly cheaper (approximately 3.32 euros per sample) than the eDNA filtration method (approximately 21.67 euros), it required considerably longer laboratory processing time. The total handling time for malacology, including snail scooping and shedding experiments, was also substantially longer (approximately two hours) than for the eDNA filtration (approximately 30 minutes to one and half hours), depending on the filtration method used (Table 8). Additionally, the malacology approach had a greater environmental impact and posed a higher health risk to people collecting snails due to prolonged water exposure, increasing the likelihood of schistosomiasis infection. In contrast, eDNA sampling required minimal direct water contact, thereby reducing risk of infection. The total processing time in the lab was also significantly shorter for the eDNA filtration method than for the conventional malacology method.

**Table 8:**
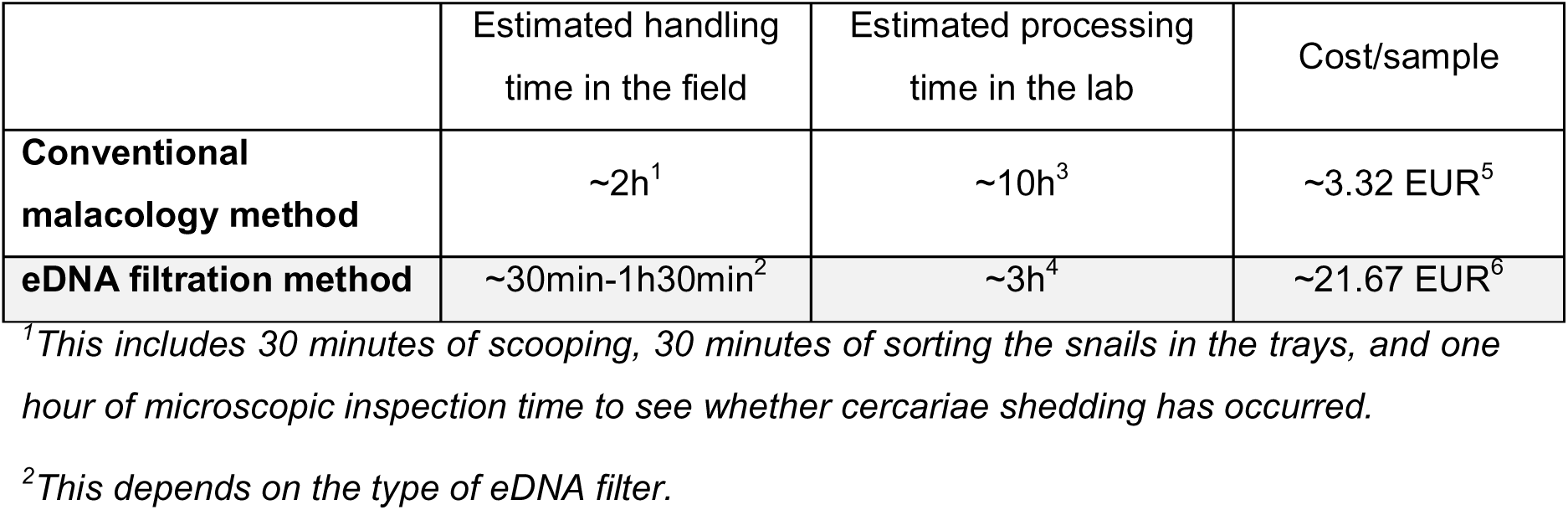
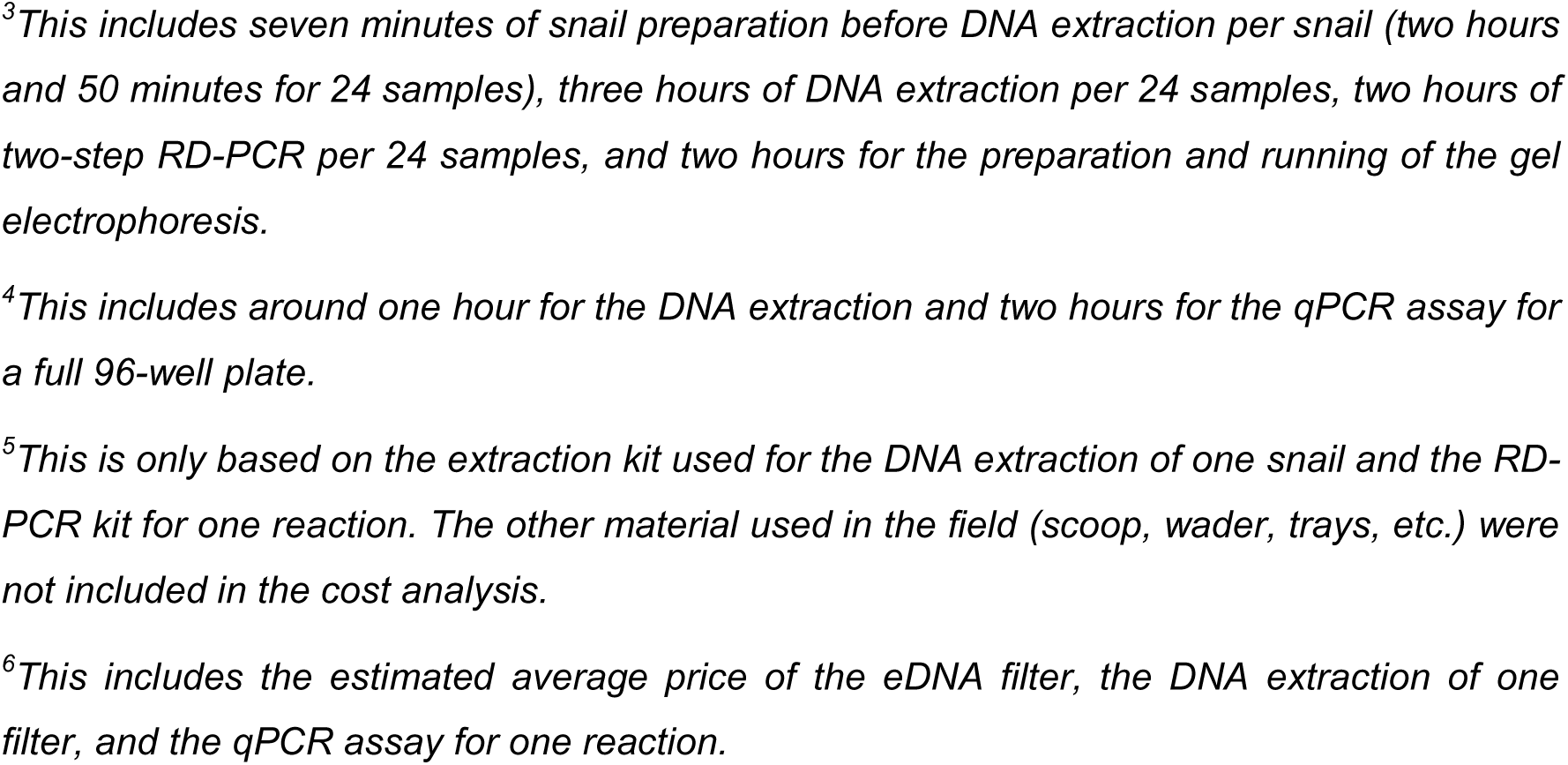
Estimated processing and handling time, and cost per sample for the eDNA filtration versus the conventional malacology method.

|  | Estimated handling time in the field | Estimated processing time in the lab | Cost/sample |
| --- | --- | --- | --- |
| **Conventional malacology method** | ~2h<sup>1</sup> | ~10h<sup>3</sup> | ~3.32 EUR<sup>5</sup> |
| **eDNA filtration method** | ~30min-1h30min<sup>2</sup> | ~3h<sup>4</sup> | ~21.67 EUR<sup>6</sup> |

<sup>1</sup>This includes 30 minutes of scooping, 30 minutes of sorting the snails in the trays, and one hour of microscopic inspection time to see whether cercariae shedding has occurred.
<sup>2</sup>This depends on the type of eDNA filter.
<sup>3</sup>This includes seven minutes of snail preparation before DNA extraction per snail (two hours and 50 minutes for 24 samples), three hours of DNA extraction per 24 samples, two hours of two-step RD-PCR per 24 samples, and two hours for the preparation and running of the gel electrophoresis.
<sup>4</sup>This includes around one hour for the DNA extraction and two hours for the qPCR assay for a full 96-well plate.
<sup>5</sup>This is only based on the extraction kit used for the DNA extraction of one snail and the RD-PCR kit for one reaction. The other material used in the field (scoop, wader, trays, etc.) were not included in the cost analysis.
<sup>6</sup>This includes the estimated average price of the eDNA filter, the DNA extraction of one filter, and the qPCR assay for one reaction.

The three eDNA filter types were compared based on their filtration rate, cost, sampling effort (defined as the time and complexity of the filtration process in the field), risk of schistosomiasis infection to the operator (duration of water exposure), and cross-contamination risk. The open membrane filter with 41 µm pore size had the highest filtration rate due to its greater mean flow rate and smaller surface area (Table 9). The two open membrane filters were significantly cheaper than other filtration types but had a higher risk of contamination and infection. In contrast, the Sylphium filters had the lowest contamination and infection risk (Table 9).

**Table 9:**
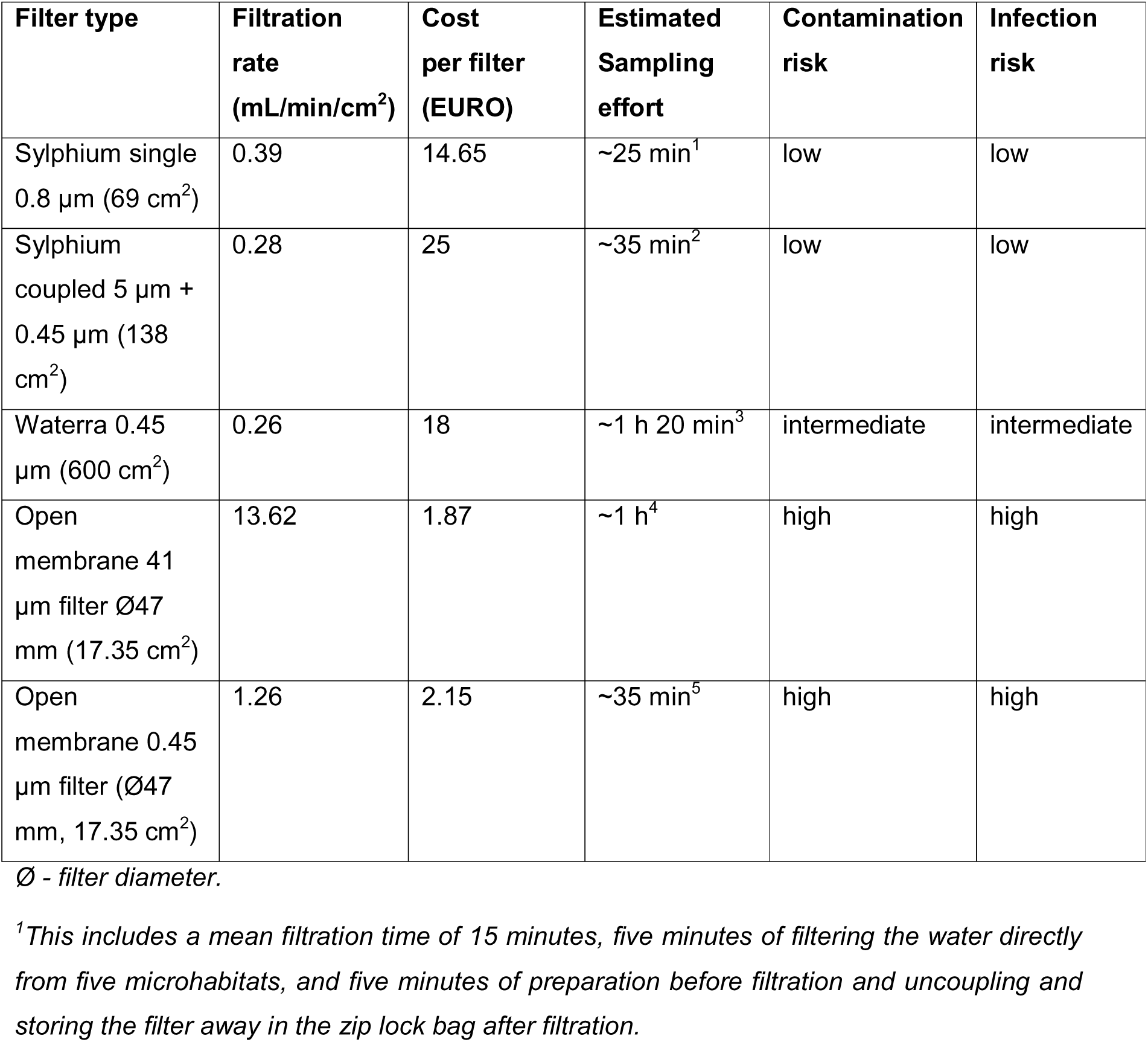
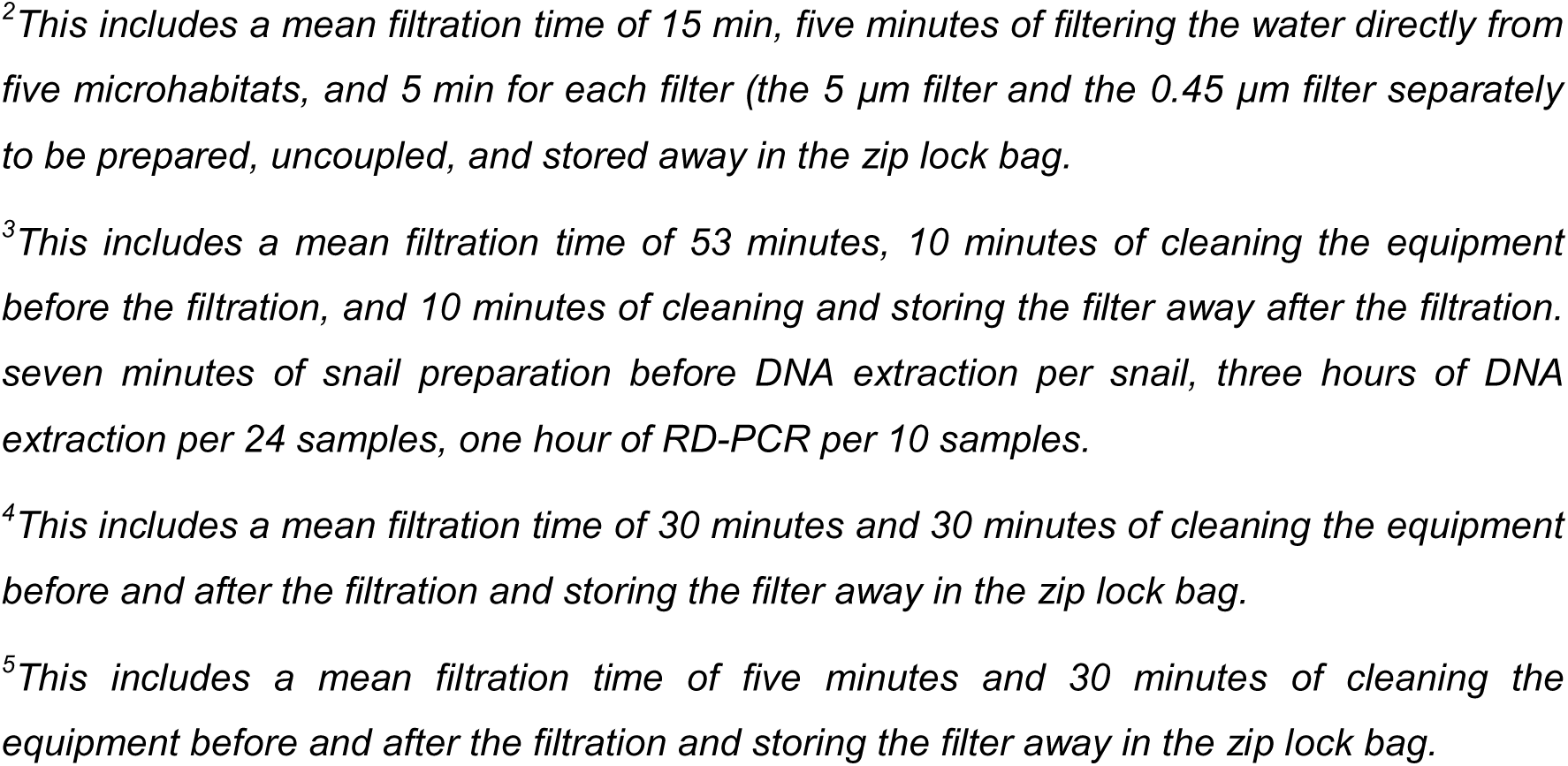
Various characteristics of the different filter types.

| Filter type | Filtration rate (mL/min/cm<sup>2</sup>) | Cost per filter (EURO) | Estimated Sampling effort | Contamination risk | Infection risk |
| --- | --- | --- | --- | --- | --- |
| Sylphium single 0.8 µm (69 cm<sup>2</sup>) | 0.39 | 14.65 | ~25 min<sup>1</sup> | low | low |
| Sylphium coupled 5 µm + 0.45 µm (138 cm<sup>2</sup>) | 0.28 | 25 | ~35 min<sup>2</sup> | low | low |
| Watterra 0.45 µm (600 cm<sup>2</sup>) | 0.26 | 18 | ~1 h 20 min<sup>3</sup> | intermediate | intermediate |
| Open membrane 41 µm filter Ø47 mm (17.35 cm<sup>2</sup>) | 13.62 | 1.87 | ~1 h<sup>4</sup> | high | high |
| Open membrane 0.45 µm filter (Ø47 mm, 17.35 cm<sup>2</sup>) | 1.26 | 2.15 | ~35 min<sup>5</sup> | high | high |

Ø - filter diameter.
<sup>1</sup>This includes a mean filtration time of 15 minutes, five minutes of filtering the water directly from five microhabitats, and five minutes of preparation before filtration and uncoupling and storing the filter away in the zip lock bag after filtration.
<sup>2</sup>This includes a mean filtration time of 15 min, five minutes of filtering the water directly from five microhabitats, and 5 min for each filter (the 5 μm filter and the 0.45 μm filter separately to be prepared, uncoupled, and stored away in the zip lock bag.
<sup>3</sup>This includes a mean filtration time of 53 minutes, 10 minutes of cleaning the equipment before the filtration, and 10 minutes of cleaning and storing the filter away after the filtration. seven minutes of snail preparation before DNA extraction per snail, three hours of DNA extraction per 24 samples, one hour of RD-PCR per 10 samples.
<sup>4</sup>This includes a mean filtration time of 30 minutes and 30 minutes of cleaning the equipment before and after the filtration and storing the filter away in the zip lock bag.
<sup>5</sup>This includes a mean filtration time of five minutes and 30 minutes of cleaning the equipment before and after the filtration and storing the filter away in the zip lock bag.

## 4. Discussion

Environmental DNA (eDNA) is increasingly recognized as a powerful tool for pathogen and vector surveillance in freshwater ecosystems (Alzaylaee et al., 2020; Kamel et al., 2021; Sato et al., 2018; Sengupta et al., 2019a, 2022). This study builds on existing research by evaluating different eDNA filtration and purification methods for detecting *S. mansoni* in endemic regions. Our findings have significant implications for optimising eDNA-based schistosomiasis surveillance and improving parasite detection sensitivity from water samples in qPCR essays.

### 4.1. The role of filtration in maximising eDNA recovery and schistosome detection

The choice of eDNA filtration method significantly influences eDNA yield. In this study, Waterra 0.45 µm capsule filters recovered the highest eDNA yields, attributed to the filter’s high-capacity of allowing larger water volumes to be processed, hence maximising eDNA retention. In contrast, Open Membrane filters yielded less DNA, likely due to faster clogging. However, despite variations in eDNA yield, *S. mansoni* detection with qPCR Ct values did not differ significantly across filtration methods. Contrastingly, positivity rates were highest with Waterra and lowest with Open Membrane. This suggests that filters enhancing eDNA recovery, such as Waterra and Sylphium filter membranes, could improve parasite detection, a hypothesis that warrants further testing with increased sampling effort.

From a practical perspective, our findings highlight a trade-off between filtration efficiency, field feasibility, and cost. Waterra and Sylphium filters require minimal handling and maximise eDNA recovery, making them ideal for large-scale surveillance. However, their cost may limit their accessibility in low-resource settings. Open membrane filters, although more affordable, pose a higher risk of contamination due to manual assembling of the filtration unit between and within sampling sites, and also high false negative rates. However, they might remain a viable alternative for fieldwork where resources are constrained if only sites with high parasite load are to be identified. Ultimately, selecting an appropriate filtration method depends on the specific study setting and the available resources.

### 4.2. Optimising eDNA purification for field samples

DNA purification efficiency significantly influenced eDNA recovery, with the Sylphium extraction method yielding the highest eDNA concentrations compared to DNeasy Blood & Tissue and ZymoBIOMICS^TM^ Miniprep kits. This is a crucial finding, as eDNA yield directly impacts detection sensitivity, particularly in low-eDNA samples where losses during purification could lead to false negatives. The superior performance of Sylphium extraction may be attributed to the differences in the extraction chemistry. The Sylphium protocol is based on DNA precipitation, where DNA is pelleted out of solution and impurities are removed through a series of wash steps. In contrast, both the DNeasy and Zymo kits use silica column-based binding, where DNA binds to a membrane and is then eluted after washing. These findings highlight that selecting an appropriate extraction method is as important as the choice of filtration. The effectiveness of eDNA-based detection depends not only on capturing target DNA from the environment but also on how well that DNA is preserved and purified during downstream processing. Future efforts in eDNA-based schistosomiasis surveillance should consider standardising purification protocols and evaluating extraction chemistry in relation to sample type, inhibitor load, and field conditions to ensure high comparability and reproducibility across studies.

### 4.3. Influence of environmental factors on eDNA persistence

Our study found no significant impact of turbidity or TDS on total eDNA yield, despite a moderate correlation between these two water quality parameters. The non-significant correlation observed between turbidity and *S. mansoni* detection (ρ = −0.359, p = 0.189) suggests that turbidity may not influence eDNA persistence. One possible explanation is that turbidity may affect eDNA transport and degradation, but its impact is site-specific and dependent on additional environmental factors. Previous studies have shown that turbidity can, in some cases, protect eDNA from degradation, as eDNA phosphate groups interact with soil surfaces to form cationic bonds (Nielsen et al., 2006; Pietramellara et al., 2009; Robson et al., 2016a). However, this adsorption is pH-dependent. Both Lake Albert and Lake Victoria are characterized by slightly alkaline pH, which decreases the strength of the electrostatic interactions with eDNA, making cationic bridging less efficient. These findings indicate that eDNA remains detectable across a range of environmental conditions, supporting its applicability in both clear and turbid habitats. Although further research is needed to assess whether extreme turbidity conditions could significantly impact *Schistosoma* eDNA degradation, dispersal, or recovery. Additional investigations incorporating factors such as pH, temperature, UV light, and microbial activity, similar to studies conducted on fish (Robson et al., 2016b) and amphibians (Buxton et al., 2017; Goldberg et al., 2018; Strickler et al., 2015) may provide a more comprehensive understanding of environmental influences on *Schistosoma* eDNA persistence.

### 4.4. eDNA versus conventional malacological surveys

The results from the eDNA qPCR amplification reinforce the effectiveness of eDNA-based monitoring for schistosomiasis surveillance. The partial agreement between eDNA detection and conventional snail surveys in Lake Albert highlights the complementary role of snail surveys in interpretating the eDNA signals. Sites with positive *Schistosoma* sp. eDNA but negative snail-based results suggest the presence of *S. mansoni* eggs or miracidia (pre-infective parasite stages), indicating environmental contamination from infected human hosts rather than active transmission. Conversely, sites with positive RD-PCR results but negative eDNA and shedding experiments may reflect prepatent snail infections, reinforcing the need for integrating multiple detection methods to accurately assess schistosomiasis active and putative transmission sites.

### 4.5. Broader implications and future directions

The successful application of eDNA in this study highlights its potential for large-scale schistosomiasis surveillance, as also demonstrated by Alzaylaee et al. (2020); Sato et al. (2018) and Sengupta et al. (2019). However, we acknowledge that our relatively small sample size limited the statistical power to robustly assess the influence of environmental factors on *S. mansoni* eDNA detectability. Future studies should incorporate more extensive spatial and temporal sampling and include a broader range of abiotic and biotic covariates known to influence eDNA occupancy and detection (Sengupta et al., 2019). Accounting for these factors would provide a more comprehensive understanding of filter performance and detection efficiency. Beyond the scope of this study, several challenges remain before widespread implementation of eDNA-based schistosomiasis surveillance is fully possible. Firstly, eDNA sampling and amplification protocols standardisation for parasite detection across varying ecological and different laboratory settings is needed to ensure comparability of results. Sampling guidelines should be developed to determine optimal water volumes, sampling frequency, and seasonal considerations to enhance detection accuracy across diverse habitats. Variations in eDNA filtration, extraction/purification, and qPCR protocols can lead to inconsistencies in detection sensitivity, which could complicate efforts to track transmission trends over time. Developing standardised guidelines for schistosomiasis eDNA surveillance, similar to those established for fish and amphibian eDNA monitoring, would greatly enhance the reliability and reproducibility of this approach (Hagerty, n.d.; Kendell et al., 2020; Melchior & Baker Cindy, 2023).

Secondly, while qPCR provides high sensitivity, the method requires expensive laboratory thermocycler equipment for DNA amplification, making it inaccessible in resource-limited settings. To address this shortcoming, eDNA-based monitoring could be coupled with point-of care isothermal DNA amplification such as Loop-Mediated Isothermal Amplification (LAMP) or Recombinase Polymerase Amplification (RPA), which are field-deployable and do not require expensive equipment (Blin et al., 2021; Fernández-Soto et al., 2014; Gandasegui et al., 2016).

Thirdly, a key limitation of eDNA is its inability to distinguish between parasite life stages. To overcome this challenge, field studies could explore the integration of environmetal RNA-based approaches to differentiate between active transmission stages (presence of infective cercariae) and potential transmission stages (presence of *Schistosoma* spp. eggs and miracidia) (Alzaylaee et al., 2020; Champion et al., 2021; Kamel et al., 2021; Sengupta et al., 2019). This strategy could enhance the specificity of eDNA monitoring by identifying life-stage-specific RNA transcripts (Cristescu, 2019; Stevens & Parsley, 2023; Yates et al., 2021), thereby providing more precise insights into ongoing transmission dynamics.

Forthly, one of the key challenges in the field was the time-consuming and impractical nature of filtering eDNA from water, particularly with open membrane and Waterra filter types. This limited the sampling to only one or two sites per day, making it less efficient than conventional malacology surveys where up to five sampling sites per day was feasible. To improve filtration, using a portable electric pump could increase efficiency, though power supply remains a concern in remote areas. Additionally, prefiltration with a larger pore size could help manage turbid water without significantly reducing eDNA capture. A larger plastic receiver bottle for open membrane filtration would also reduce sampling effort and contamination risks by minimising the need to frequently disconnect and empty the collection bottle. Further improvements include refining filter preservation methods to prevent eDNA loss, such as inserting a cotton ball between the filter and silica beads or using a paper coin envelope system. Collecting water without entering the lake could also reduce infection risks, potentially through the use of boats, an extendable rod, or peristaltic pumps, though field conditions may limit these solutions. Future studies should optimise water volume filtration across varying turbidity levels to balance efficiency and eDNA yield while minimising costs and workload.

Lastly, understanding the ecological and hydrological dynamics of *Schistosoma* spp. eDNA in freshwater systems will be critical for optimising sampling strategies. Factors such as eDNA release by the parasite, transport, degradation rates, and the influence of seasonal fluctuations on eDNA detectability, require further investigation to refine predictive models for disease transmission risk. Addressing these challenges will be essential for harnessing eDNA as a reliable and scalable tool for schistosomiasis surveillance in endemic regions.

## 6. Conclusion

This study demonstrates the utility of eDNA as a tool for schistosomiasis surveillance, while highlighting differences in efficiency and sensitivity of three eDNA filtration and purification methods from water samples. While Waterra filters and Sylphium extraction maximised eDNA recovery, our results suggest that lower-yielding filtration approaches such as Open membrane filtration, can still provide sufficient eDNA for *S. mansoni* detection in sites with high parasite loads. This makes eDNA a versatile and scalable tool schistosomiasis monitoring, adaptable to different resource and transmission settings. However, to fully integrate eDNA into schistosomiasis control programs, future efforts should focus on protocol standardisation, validation across diverse ecological settings, and the development of field-deployable isothermal amplification strategies such as LAMP suitable for low-resource settings. As eDNA surveillance advances, it has the potential to revolutionise parasitic disease ecoepidemiology, offering a non-invasive, efficient, and scalable solution for global schistosomiasis control efforts.

## CRediT authorship contribution statement

**Cecilia Wangari Wambui:** Writing - original draft and final copy, Writing - review & editing Methodology, Formal analysis, Conceptualisation.

**Mila Viaene:** Writing - original draft and final copy, Writing - review & editing, Data collection, Methodology, Formal analysis, Conceptualisation.

**Hannah Njiriku Mwangi:** Writing - review & editing, Data collection, Methodology, Conceptualisation.

**Benjamin André:** Writing - review & editing, Data collection, Conceptualisation.

**David Were Oguttu:** Writing - review & editing, Sampling sites selection, Fieldwork coordination.

**Casim Umba Tolo:** Sampling sites selection, Project administration and coordination.

**Bart Hellemans:** Writing - review & editing, Methodology, Formal analysis.

**Tine Huyse:** Writing - review & editing, Conceptualisation, Supervision, Securing funding, Project administration and coordination.

**Hugo F. Gante:**Writing - review & editing, Methodology, Formal analysis, Conceptualisation, Supervision, Securing funding, Project administration and coordination.

## Declaration of competing interest

All authors declare no competing financial interests.

## Acknowledgements

We appreciate the U-SMRC project of the Uganda Virus Research Institute (UVRI) for selecting study sites and signing Prior informed consent (PIC) and mutually agreed terms for eDNA studies with local communities and cherishing good collaboration. We acknowledge UVRI for Material Transfer Agreements (MTAs) and UNCST for granting permission to export samples from Uganda to Belgium. We also thank the Natural History Museum providing Schistosoma adult worms specimen. We thank field assistants and community guides for support during data collection.

## Funding

This research was supported by: (i) KU Leuven Research Fund grant STG/21/044 to HFG and the American Leprosy Mission 2023 NTD Innovation Prize (USA295-NTD Innovation Prize FY24), awarded to the ATRAP (Action Towards Reducing Aquatic snail-borne Parasitic Disease) project for providing fieldwork sampling material, laboratory reagents and consumables. (ii) The VLIR-UOS travel grant for funding the travel expenses to Uganda and the U-SMRC project (USA-NIH grant number U01Al168609).

---

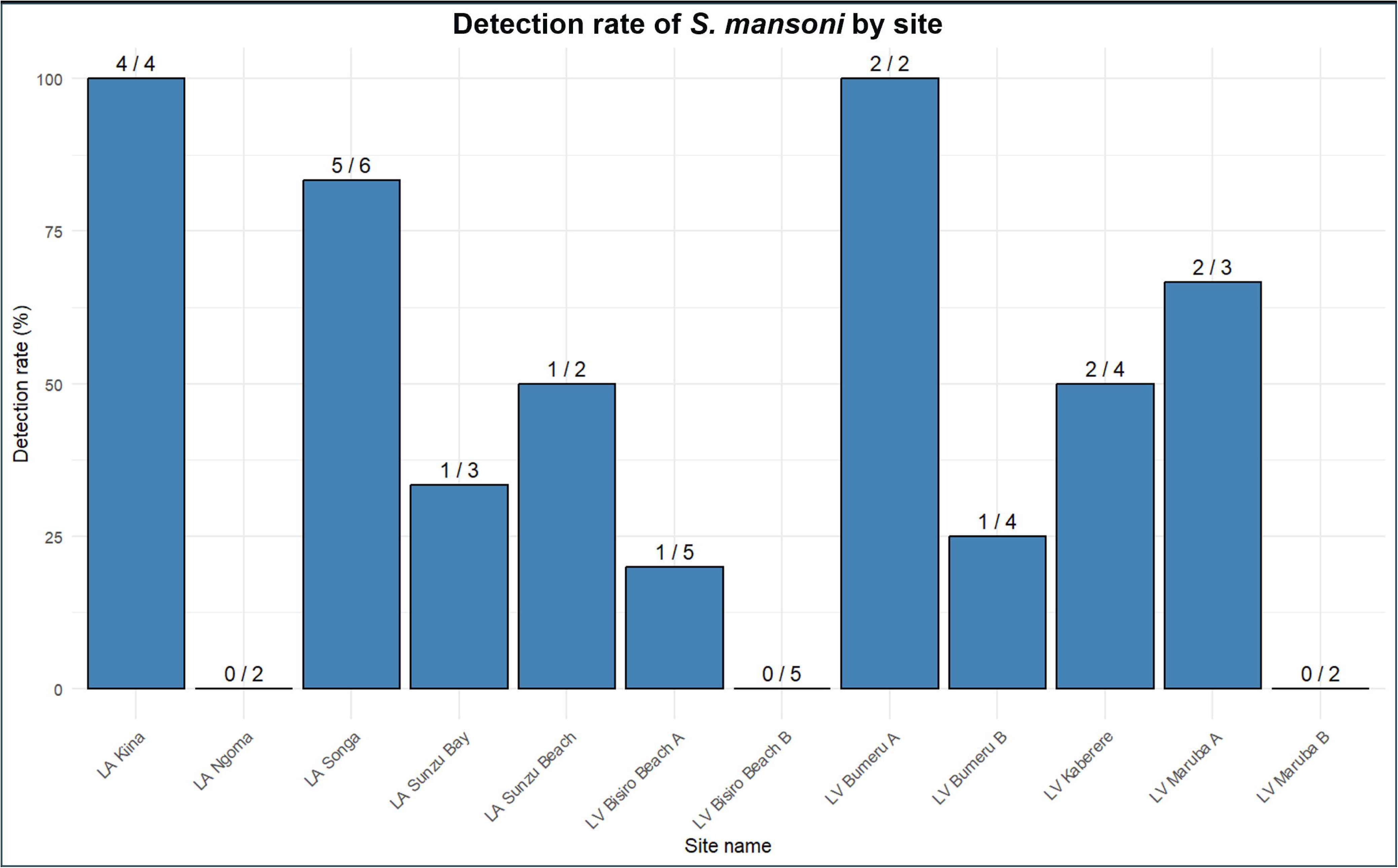

## Notes

### Competing Interest Statement

The authors have declared no competing interest.

