## Supplementary information for "Improving eDNA filtration and purification for qPCR detection of *Schistosoma mansoni* in lakes Albert and Victoria"

Table 1: The accession numbers of 50 trematode mitogenomes (adapted from (Douchet et al., 2022)) and 37 COI gene sequences of different *Schistosoma* isolates representing different geographical regions of Africa used for *in silico* testing.

| **Accession number** | **Species** | **Accession number** | **Species** |
| --- | --- | --- | --- |
| NC_029757.1 | *Brachycladium goliath* | NC_039532.1 | *Echinostoma miyagawai* |
| NC_044643.1 | *Postharmostomum commutatum* | NC_037150.1 | *Artyfechinostomum sufrartyfex* |
| NC_027082.1 | *Clinostomum complanatum* | KM111525.1 | *Hypoderaeum sp.* |
| NC_039780.1 | *Cyathocotyle prussica* | MN496162.1 | *Echinostoma revolutum* |
| NC_044135.1 | *Tracheophilus cymbius* | MK238506.1 | *Amphimerus sp.* |
| NC_042722.1 | *Uvitellina sp.* | NC_023095.1 | *Paramphistomum cervi* |
| NC_026916.1 | *Eurytrema pancreaticum* | NC_027271.1 | *Calicophoron microbothrioides* |
| NC_025280.1 | *Dicrocoelium dendriticum* | NC_028071.1 | *Orthocoelium streptocoelium* |
| NC_002546.1 | *Fasciola hepatica* | NC_027958.1 | *Explanatum explanatum* |
| NC_030528.1 | *Fasciolopsis buski* | NC_042482.1 | *Plagiorchis maculosus* |
| NC_024025.1 | *Fasciola gigantica* | NC_036411.1 | *Trichobilharzia szidati* |
| NC_029481.1 | *Fascioloides magna* | NC_009680.1 | *Trichobilharzia regenti* |
| KX787886.1 | *Fasciola jacksoni* | NC_008074.1 | *Schistosoma haematobium* |
| NC_030530.1 | *Homalogaster paloniae* | NC_008067.1 | *Schistosoma spindale* |
| NC_028001.1 | *Fischoederius elongatus* | NC_027673.1 | *Paragonimus sp.* |
| NC_030529.1 | *Fischoederius cobboldi* | NC_039430.1 | *Paragonimus heterotremus* |
| NC_027833.1 | *Gastrothylax crumenifer* | NC_032032.1 | *Paragonimus ohirai* |
| NC_022433.1 | *Haplorchis taichui* | NC_002354.2 | *Paragonimus westermani* |
| NC_023249.1 | *Metagonimus yokogawai* | NC_002529.1 | *Schistosoma mekongi* |
| MG792058.1 | *Acanthoparyphium sp.* | NC_002545.1 | *Schistosoma mansoni* |
| NC_027112.1 | *Ogmocotyle sikae* | OX104043.1 | *Schistosoma rodhaini* |
| KR006935.1 | *Ogmocotyle sp.* | OX103949.1 | *Schistosoma mattheei* |
| NC_012147.2 | *Clonorchis sinensis* | OQ568668.1 | *Schistosoma haematobium* |
| NC_028008.1 | *Metorchis orientalis* | OX103960.1 | *Schistosoma bovis* |
| NC_011127.2 | *Opisthorchis felineus* | OX103896.1 | *Schistosoma guineensis* |
| NC_025279.1 | *Dicrocoelium chinensis* | OX104147.1 | *Schistosoma curassoni* |
| NC_030518.1 | *Echinochasmus japonicus* | OX104030.1 | *Schistosoma margrebowiei* |
| MH212284.1 | *Echinostoma sp.* | KY967520.1 | *Schistosoma haematobium* |
| NC_028010.1 | *Echinostoma hortense* | KU196388.1 | *Schistosoma japonicum* |
| HQ839768.1 | *Schistosoma mansoni* | HQ122388.1 | *Schistosoma mansoni* |
| GU294837.1 | *Schistosoma mansoni* | LN876761.1 | *Schistosoma mansoni* |
| KT354659.1 | *Schistosoma haematobium* | KP343672.1 | *Schistosoma mansoni* |
| KP343649.1 | *Schistosoma mansoni* | GU294793.1 | *Schistosoma bovis* |
| OX103960.1 | *Schistosoma bovis* | OZ022085.1 | *Schistosoma bovis* |
| AY157212.1 | *Schistosoma bovis* | OZ022082.1 | *Schistosoma bovis* |
| MH014043.1 | *Schistosoma bovis* | MF919407.1 | *Schistosoma bovis* |
| AY157210.1 | *Schistosoma currasoni* | OZ022086.1 | *Schistosoma currasoni* |
| OZ022039.1 | *Schistosoma bovis x Schistosoma haematobium* | OZ022196.1 | *Schistosoma bovis x Schistosoma currasoni* |
| DQ354364.1 | *Schistosoma guineensis x Schistosoma intercalatum* | JQ082121.1 | *Schistosoma haematobium* |
| MT886703.1 | *Schistosoma haematobium* | OP235429.1 | *Schistosoma haematobium* |
| OP779237.1 | *Schistosoma haematobium* | OL840258.1 | *Schistosoma haematobium* |
| JQ397397.1 | *Schistosoma haematobium* | GU257397.1 | *Schistosoma haematobium* |
| OK314869.1 | *Schistosoma haematobium* | MT380523.1 | *Schistosoma haematobium* |
| JQ397353.1 | *Schistosoma haematobium* |  |  |

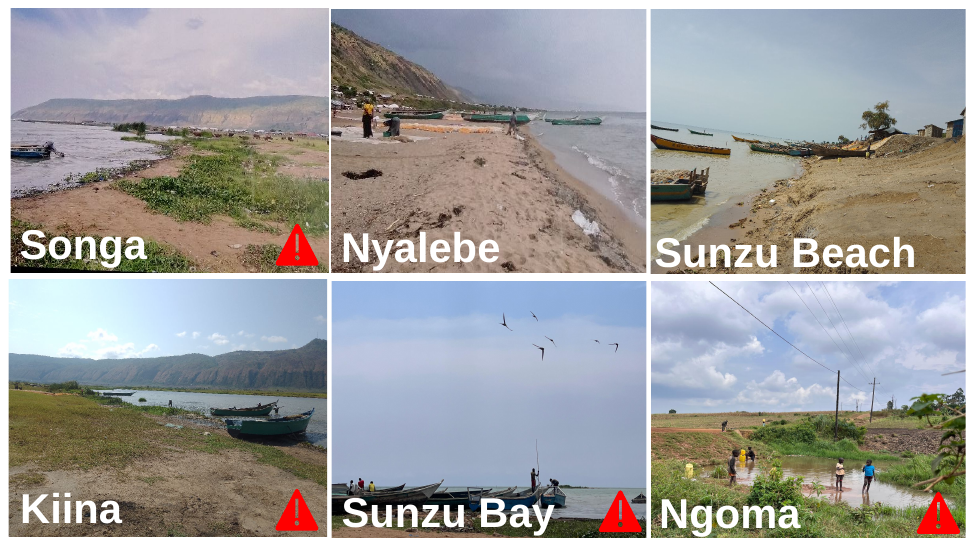

**Figure 1: Pictures of the different sampling sites along Lake Albert.**

**Figure 2: Pictures of the different sampling sites along Lake Victoria.**

**Table 2: Overview of the sampling sites with their characteristics**

| *Site* | *Site name* | *Date* | *Coordinates* | *weather* | *Site type* | *Substrate* | *activities* | *surrounding* | *Environment modification* |
| --- | --- | --- | --- | --- | --- | --- | --- | --- | --- |
| *LA01* | *Kiina* | *15/08/2023* | N 1°12'23''  E 30°43'16'' | *Sunny* | *Lake beach* | *Mud, sand* | *Washing items, fishing, watering animals, drinking* | *Grazing* | *Vegetation clearance* |
| *LA02* | *Sunzu Bay* | *16/08/2023* | N 1°13'19"  E 30°43'30'' | *NA* | *Lake beach* | *Mud, sand* | *Collecting water, swimming, child play, washing items, fishing* | *Grazing* | *Vegetation clearance* |
| *LA03* | *Sunzu Beach* | *16/08/2023* | N 1°13'57"  E 30°43'57'' | *Sunny* | *Lake beach* | *Sand* | *Collecting water, swimming, child play, washing items, fishing* | *Housing* | *Vegetation clearance* |
| *LA04* | *Songa* | *17/08/2023* | N 1°14'27"  E 30°44'8'' | *Sunny windy* | *Lake beach* | *Mud, loose rock, sand* | *Collecting water, swimming, child play, washing items, watering animals* | *Grazing* | *Vegetation clearance* |
| *LA05* | *Nyalebe* | *17/08/2023* | N 1° 16'36''  E 30°45'32'' | *Rainy* | *Lake beach* | *Sand* | *Collecting water, swimming, child play, washing items, fishing, watering animals* | *Housing* | *Vegetation clearance* |
| *LA06* | *Ngoma* | *18/09/2023* | N 1°13'27"  E 30°46'41" | *Cloudy* | *Puddle* | *mud, rocky stone, clay* | *Collecting water, swimming, child play, washing items, watering animals* | *Agriculture* | *Vegetation clearance* |
| *LA07* | *Nyampindu* | *18/09/2023* | N 1°10'15"  E 30°44'15" | *Sunny windy* | *Puddle* | *Mud, clay* | *Collecting water, washing items, watering animals* | *Agriculture, grazing* | *NA* |
| *LV01A* | *Bisiro Beach* | *15/09/2023* | N 0°11'11''  E 33°51'10'' | *Sunny* | *Lake beach* | *Mud, rocky stone, sand* | *Collecting water, batching, washing items, fishing* | *Housing, grazing* | *Vegetation clearance* |
| *LV01B* | *Bisiro Beach* | *15/09/2023* | N 0°11'03''  E 33°51'21'' | *Cloudy* | *Lake marsh* | *Mud* | *Collecting water, watering animals* | *Agriculture, grazing* | *No modification* |
| *LV02A* | *Buduma* | *16/09/2023* | N 0°8'41''  E 33°48'30'' | *Sunny* | *Lake beach* | *Rocky stone, loose rock, sand* | *Collecting water, bathing, washing items, fishing* | *Housing* | *Vegetation clearance* |
| *LV02B* | *Buduma* | *16/09/2023* | N 0°08'55"  E 33°48'37" | *Sunny* | *Lake marsh* | *Mud, rocky stone* | *Collecting water, bathing, swimming, child play, washing items, watering animals* | *Grazing* | *Vegetation clearance* |
| *LV03A* | *Bumeru* | *17/09/2023* | N 0°12'37''  E 33°42'36'' | *Sunny* | *Lake beach* | *Mud, clay* | *Collecting water, bathing, washing items, fishing, watering animals* | *Grazing* | *Vegetation clearance* |
| *LV03B* | *Bumeru* | *17/09/2023* | N 0°12'18''  E 33°42'18'' | *Sunny* | *Lake marsh* | *Mud, clay* | *Collecting water, swimming, child play, washing items, fishing* | *Agriculture, grazing* | *Vegetation clearance* |
| *LV04A* | *Maruba* | *18/09/2023* | N 0°16'9''  E 33°41'28'' | *Very cloudy* | *Lake beach/marsh* | *Mud, sand* | *collecting water, washing items, fishing, watering animals* | *Agriculture, grazing* | *NA* |
| *LV04B* | *Maruba* | *18/09/2023* | N 0°15'53''  E 33°41'30'' | *Cloudy* | *Lake marsh* | *Mud, sand* | *Collecting water, washing items, fishing, watering animals* | *Agriculture, grazing* | *Vegetation clearance* |
| *LV05* | *Kaberere* | *19/09/2023* | N 0°30'08"  E 33°52'32'' | *Sunny* | *Lake marsh rice field* | *Mud, rocky stone, loose rock, sand* | *Collecting water, fishing, watering animals* | *Grazing* | *Vegetation clearance* |
| *LV06A* | *Lugala* | *20/09/2023* | N 0°11'57.2"  E 33°54'20" | *Sunny* | *Lake beach* | *Sand* | *Collecting water, bathing, washing items, fishing, watering animals* | *Agriculture, grazing, drying fish* | *Vegetation clearance* |
| *LV06B* | *Lugala* | *20/09/2023* | N 0°11'52.9"  E 33°54'47" | *Rainy* | *Lake marsh* | *Mud, sand* | *Collecting water, washing items* | *Grazing* | *No modification* |

**Table 3: Abiotic water parameters measured per sampling site. Abbreviations: conductivity (Conduct.), water temperature (Water T), dissolved oxygen (D.O), and total dissolved solids (TDS).**

| **Site** | **pH** | **Conduct.**  **(µS/cm)** | **Water T**  **(°C)** | **D.O**  **(mg/L)** | | **D.O (%)** | **TDS (ppm)** | **Salinity (PSU)** | **Turbidity (FNU)** | **Resistivity (MΩ.cm)** | | | **Nitrate (mg/L)** | **Phosphate (mg/L)** |
| --- | --- | --- | --- | --- | --- | --- | --- | --- | --- | --- | --- | --- | --- | --- |
| **LA01** | 8.7 | 581 | 29.1 | 294 | | NA | NA | NA | NA | NA | | | 0 | 2.5 |
| **LA02** | 8.79 | 593 | 30.54 | 4.23 | | 60.5 | 290 | 0.28 | 69 | 0.0017 | | | 0 | 2.5 |
| **LA­03** | 9.17 | 569 | 31.99 | 4.13 | | 60.3 | 297 | 0.28 | 123 | 0.0017 | | | 0 | 2.5 |
| **LA04** | 9.24 | 566 | 27.68 | 4.8 | | 64.8 | 285 | 0.27 | 47.6 | 0.0018 | | | 133 | 0.57 |
| **LA05** | 8.64 | 226 | 28.99 | 4.64 | | 64.7 | 283 | 0.27 | 15.9 | 0.0018 | | | 0 | 2.5 |
| **LA06** | 7.91 | 298 | 21.99 | 7.53 | | 97.7 | 113 | 0.11 | 86.7 | 0.0041 | | | 0 | 0.82 |
| **LA07** | 8.67 | 568 | 24.52 | 4.75 | | 64.3 | 149 | 0.14 | 218 | 0.0034 | | | 0 | 2.5 |
| **LV01A** | 9.73 | 105 | 30.27 | 0.11 | | 1.7 | 53 | 0.05 | 15.3 | 0.0095 | | | 0 | 0.38 |
| **LV01B** | 9.58 | 180 | 27.65 | 0.22 | | 3.2 | 90 | 0.08 | 7.1 | 0.0056 | | | 0 | 2.5 |
| **LV02A** | 9.29 | 106 | 28.35 | 0.46 | | 6.7 | 53 | 0.05 | 13.7 | 0.009 | | | 0 | 2.5 |
| **LV02B** | 8.01 | 107 | 29.61 | 0.61 | | 9.1 | 54 | 0.05 | 34.6 | 0.0094 | | | 5.6 | 2.5 |
| **LV03A** | 8.68 | 116 | 27.96 | 1.27 | | 18.9 | 57 | 0.05 | 19.1 | 0.0086 | | | 0 | 2.5 |
| **LV03B** | 8.57 | 166 | 30.47 | 0.88 | | 13.5 | 83 | 0.08 | 42.5 | 0.006 | | | 5.1 | 2.5 |
| **LV04A** | 8.46 | 106 | 26.22 | 1.68 | | 24 | 53 | 0.05 | 39.4 | 0.0094 | | | 0 | 2.5 |
| **LV04B** | 8.34 | 118 | 27.87 | 1.38 | | 20.4 | 56 | 0.05 | 220 | 0.0089 | | | 0 | 2.5 |
| **LV05A** | 8.68 | 402 | 26.53 | 1.28 | | 18.1 | 201 | 0.19 | 23.3 | 0.0025 | | | 0 | 2.5 |
| **LV06A** | 8.49 | 128 | 28.13 | 0.7 | 10.3 | | 55 | 0.05 | 5.1 | | 0.0092 | 0 | | 2.5 |
| **LV06B** | 8.53 | 589 | 28.84 | 0.55 | 8.1 | | 59 | 0.05 | 81 | | 0.0084 | 0 | | 2.5 |

**Table 4: Overview of the biotic and social characteristics measured at each sampling site**

| *Site* | *Water level* | *Water turbidity* | *Wave action* | *Wave strength* | *Animals* | *type vegetation* | *# men* | *# women* | *# child* | *Altitude* |
| --- | --- | --- | --- | --- | --- | --- | --- | --- | --- | --- |
| *LA01* | *Low* | *Low* | *Yes* | *Weak* | *Cattle, goat* | *Submerged, emerged, floating, rooted* | 5 | 1 | 1 | 613.45 |
| *LA02* | *Low* | *Medium* | *Yes* | *Weak* | *Cattle, goat* | *Submerged, emerged, floating, rooted* | 6 | 2 | 8 | 601.89 |
| *LA03* | *Moderate* | *Medium* | *Yes* | *Weak* | *Cattle, goat* | *Submerged, emerged, floating, rooted* | 11 | 9 | 5 | 602.69 |
| *LA04* | *Moderate* | *Medium* | *Yes* | *Strong* | *Cattle, goat* | *Submerged, emerged, floating, rooted* | 13 | 5 | 2 | 606.76 |
| *LA05* | *Moderate* | *Medium* | *Yes* | *Strong* | *No animals* | *Submerged, floating* | 8 | 4 | 9 | 615 |
| *LA06* | *Moderate* | *Medium* | *No* | *NA* | *Goat* | *Emerged, floating, rooted* | 0 | 3 | 3 | 1084.78 |
| *LA07* | *High* | *High* | *No* | *NA* | *Goat* | *Emerged, floating, rooted* | 0 | 0 | 0 | 1141.02 |
| *LV01A* | *High* | *Medium* | *Yes* | *Weak* | *Cattle,*  *goat, duck, chicken, other* | *Emerged, floating,*  *rooted, other* | 2 | 5 | *5* | 1124.83 |
| *LV01B* | *Low* | *High* | *No* | *NA* | *Cattle* | *Submerged,*  *floating, rooted* | 0 | 0 | *1* | 1130 |
| *LV02A* | *NA* | *Low* | *No* | *NA* | *No animals* | *Floating, rooted* | 6 | 3 | *1* | 1122.73 |
| *LV02B* | *Moderate* | *Medium* | *Yes* | *Normal* | *Cattle* | *Submerged,*  *emerged, floating, rooted* | 0 | 3 | *3* | 1136.83 |
| *LV03A* | *low* | *High* | *No* | *NA* | *Cattle, goat, duck, chicken, dog* | *Submerged,*  *emerged, floating, rooted* | 7 | 8 | *8* | 1181.54 |
| *LV03B* | *Low* | *High* | *Yes* | *Normal* | *Dog, other* | *Submerged,*  *emerged, rooted* | 2 | 3 | *5* | 1121.41 |
| *LV04A* | *Low* | *Medium* | *Yes* | *Normal* | *Cattle, goat, chicken* | *Submerged,*  *emerged, floating, rooted* | 3 | 4 | *3* | 1129.42 |
| *LV04B* | *Low* | *High* | *Yes* | *Normal* | *Cattle, goat* | *Submerged,*  *emerged, floating, rooted* | 1 | 0 | 1 | 1137.96 |
| *LV05A* | *Moderate* | *Medium* | *No* | *Normal* | *Cattle, dog* | *Submerged,*  *emerged, floating, rooted* | 2 | 0 | 0 | 1136.83 |
| *LV06A* | *low* | *Low* | *No* | *NA* | *Cattle, goat, other* | *Submerged, emerged, rooted* | 3 | 11 | 5 | 1134.96 |
| *LV06B* | *Low* | *High* | *No* | *NA* | *Cattle, goat* | *Submerged, emerged, rooted* | 0 | 1 | 2 | 1101.46 |

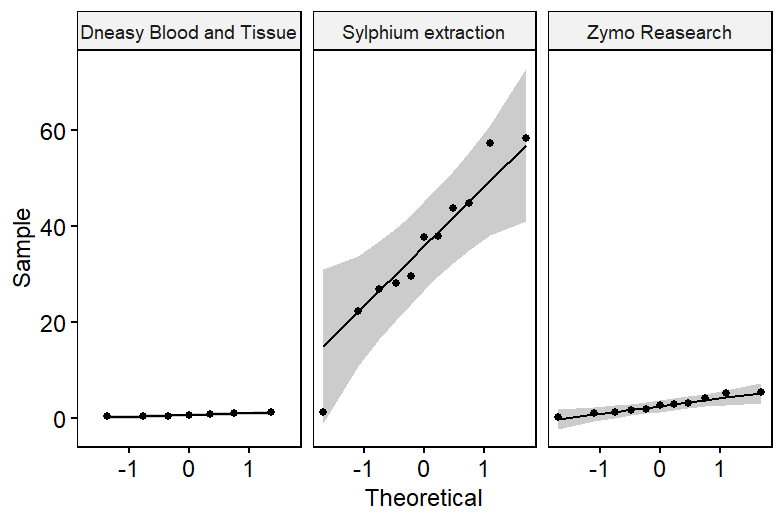

Figure 3: Normality test for the three purification methods: DNeasy Blood and Tissue, Sylphium extraction and Zymo Research eDNA yield data

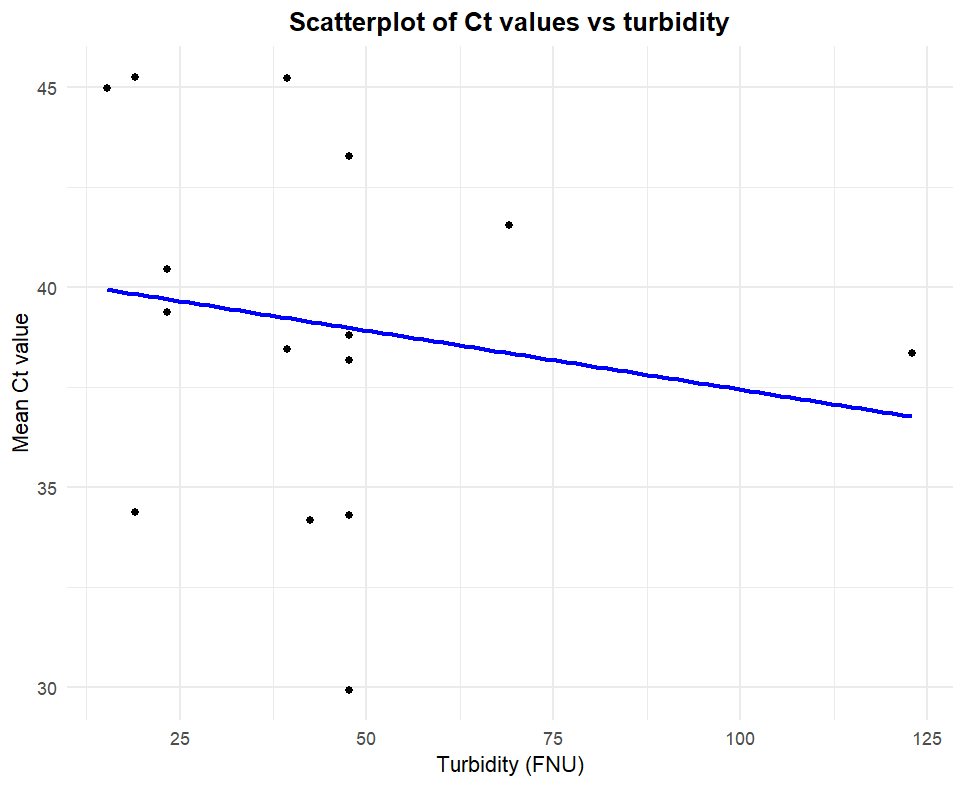

Figure 4: Scatterplot showing the relationship between mean Ct values and water turbidity (FNU). The blue line represents the fitted linear regression model.
